# Accurate protein stability prediction for small domains using mega-scale experiments

**DOI:** 10.64898/2026.05.19.726285

**Authors:** Yehlin Cho, Kotaro Tsuboyama, Theodore J. Litberg, Michelle D. Jung, Adunoluwa Obisesan, Qian Wang, Claire M. Phoumyvong, Jane Thibeault, Sergey Ovchinnikov, Gabriel J. Rocklin

## Abstract

Predicting absolute protein folding stability is a long-standing challenge in biophysics, with broad applications in protein design and in understanding genetic variation and evolution. Physics-based simulations have shown limited success at predicting stability and are often computationally intractable, and machine learning methods have been constrained by the lack of sufficiently large experimental datasets. We recently introduced cDNA display proteolysis, a cell-free approach that can measure folding stability for nearly one million protein domains in parallel. Here, we applied this method to measure stability for 1.8 million diverse protein domains 60-80 amino acids in length primarily taken from the MGnify metagenomic database and spanning over 200,000 sequence families. Using this new “MGnify Stability dataset”, we developed the predictive models SaProtΔG and ESM3ΔG, which accurately predict absolute folding stability for small domains with root mean squared error of 0.8 kcal/mol over a 6 kcal/mol range (Spearman rank correlation of 0.88). These predictors show high accuracy at predicting effects of substitutions, insertions, and deletions, successfully identify global trends toward higher stability in thermophilic organisms, and improve discrimination of stable and unstable computationally designed proteins. Our results illustrate how megascale biophysical measurements can complement existing evolutionary and structural data to enable accurate absolute stability prediction for small domains.

## Introduction

Predicting protein folding stability (the ratio of folded to unfolded molecules at equilibrium for a specific protein sequence) remains one of the oldest unsolved problems in molecular biophysics (Richards 1997). Over 50 years ago, researchers recognized that different proteins varied widely in stability (Pace and Alexander 1971), yet no simple physicochemical properties (Goldsack 1970; Singleton and Amelunxen 1973) or structural features (Matthews et al. 1974; Grütter et al. 1979) could be found that clearly and universally differentiated higher and lower stability proteins. This complexity was reinforced by protein design studies that unintentionally produced numerous protein-like amino acid sequences (Kamtekar et al. 1993) with plausible three-dimensional structures (Koga et al. 2012; Rocklin et al. 2017; Koepnick et al. 2019; Kim et al. 2022; Haddox et al. 2025; Cho et al. 2025; Dantas et al. 2003) that lacked any observable folding stability (Pan and Kortemme 2021; Garcia et al. 2026). The fundamental difficulty is that stability reflects the balance of many large, opposing energetic terms: folding leads to a large loss of conformational entropy and protein-solvent interactions (Clark et al. 2020) that must be overcome by favorable intra-protein interactions including hydrophobic burial, van der Waals interactions, hydrogen bonding, and electrostatic interactions (Kauzmann 1959; Dill 1990b; Privalov 1990). These large opposing energies (in excess of 100 kcal/mol, (Schellman 1997)) result in marginal stabilities typically below 6 kcal/mol (Walker et al. 2019)—on the order of just a few interactions (Nick Pace et al. 2014). As a result, every interaction can play a pivotal role in folding stability (memorably described by Petsko: “So is EVERYTHING Important EVERYWHERE?” (Petsko 2001)). Despite this complexity, understanding and predicting folding stability remains centrally important due to the critical role stability plays across scientific areas, including in human disease (Beltran et al. 2025; Dobson 2003), protein evolution (Zeldovich et al. 2007; Bloom et al. 2006), therapeutic development (Roberts 2014; Rahban et al. 2023), and biotechnology (Sumida et al. 2024; Stimple et al. 2020; Romero and Arnold 2009). Furthermore, because the same biophysical forces that drive folding also drive protein-protein interactions (Almeida et al. 2021; Deng et al. 2025), protein-drug binding (Myslinski et al. 2011; Klebe 2015), conformational change, and allostery (Radley et al. 2003), accurate measurements and models of stability may provide a route to illuminate these areas as well.

Several other challenges have limited the development of accurate models to predict stability. First, folding stability depends on the full ensemble of both folded and unfolded conformations (Shortle 1996; Dill and MacCallum 2012), including the highly heterogeneous unfolded ensemble, which is a difficult modeling problem (Lee et al. 2026; Cho et al. 2025). To mitigate the need to model the full structure and unfolded ensemble, most predictive models for folding stability (Dieckhaus et al. 2024; Li and Luo 2025) are limited to computing localized effects of point mutations (ΔΔG) instead of the overall folding stability (ΔG), though this substantially limits the utility of these models, especially for *de novo* protein design. Second, folding stability is sensitive to environmental conditions including pH, temperature, and ionic strength (Pace et al. 1996; Kumar and Nussinov 2002). This environmental sensitivity makes it difficult to derive accurate stability models from databases of stability measurements performed under different experimental conditions (Nikam et al. 2021; Xavier et al. 2021). Third, many of the driving forces contributing to folding stability (such as the hydrophobic effect and electrostatic interactions in partially polarizable environments) are fundamentally challenging to model and context dependent (Zhou and Pang 2018; Pace et al. 1996; Lins and Brasseur 1995).

Computational approaches have not yet overcome the challenges of predicting stability. Physics-based simulations using molecular dynamics can, in principle, estimate ΔG directly, but this requires extensive sampling of both folded and unfolded ensembles – an enormous computational challenge for proteins larger than a few dozen residues. The accuracy of simulations also remains limited by approximate force fields that cannot yet be meaningfully validated for folding stability prediction due to the computational expense (Karplus and McCammon 2002; Shirts and Pande 2007; Piana et al. 2014; Robustelli et al. 2018). Machine learning approaches hold promise for predicting folding stability, but these have been constrained by the availability of experimental data. Traditionally, measuring folding stability required purifying each individual protein (Pace and Alexander 1971), and the total number of stability measurements (thousands of measurements from several hundred protein folds (Xavier et al. 2021; Kumar et al. 2006)) never proved sufficient for developing accurate machine learning models to predict ΔG. To address the lack of large, high-quality protein stability data, we recently developed the cDNA display proteolysis method for measuring folding stability and constructed the “Megascale” dataset of 776,000 stability measurements of mutant proteins taken from 479 wild-type protein domains (Tsuboyama et al. 2023). These unique large-scale data have substantially improved ΔΔG prediction (Dieckhaus et al. 2024; Chen et al. 2025; Li and Luo 2025) and have also enabled new progress at predicting ΔG (Lewis et al. 2025; Lee et al. 2026). However, the limited structural diversity of the Megascale dataset is insufficient to capture the full diversity of protein domain architectures and likely too sparse to train accurate, generalizable predictors of ΔG.

Here, we used cDNA display proteolysis (Tsuboyama et al. 2023) to experimentally measure folding stability for nearly two million protein domains (Fig. 1a) from the MGnify metagenomic database (Richardson et al. 2023), and used these data to develop accurate models to predict folding stability (ΔG_unfold_) for domains under 80 amino acids in length. The domains in our new MGnify Stability dataset are vastly more diverse than existing stability datasets. These domains form 211,020 sequence clusters at 30% sequence identity (compared to 199 in the original Megascale dataset or 470 wild-type proteins in ThermoMutDB (Xavier et al. 2021)) and form 129,303 structural clusters with Tm-score < 0.5 (Fig. 1b). Using this new MGnify Stability dataset, we fine-tuned the structure-aware protein language models SaProt (Su et al. 2023) and ESM3 (Hayes et al. 2025) to develop the predictive models SaProtΔG and ESM3ΔG. These models take a protein domain’s sequence and predicted structure as input to predict its absolute folding stability under the conditions of our experiment (pH 7.4, 298K). Both models achieve strong performance on held-out test data with limited sequence or structural similarity to the training data. Although these models were fine-tuned using our data from domains <80 amino acids in length, both models also show promising performance on larger proteins. This includes predicting mutant effects in larger domains, distinguishing mesophilic and thermophilic protein sequences, ranking nanobodies by apparent melting temperature, and discriminating between active versus inactive computationally designed binders. Collectively, these results demonstrate that large-scale, uniform measurements combined with structure-aware modeling open a new path toward solving the long-standing stability prediction problem.

**Figure 1.**
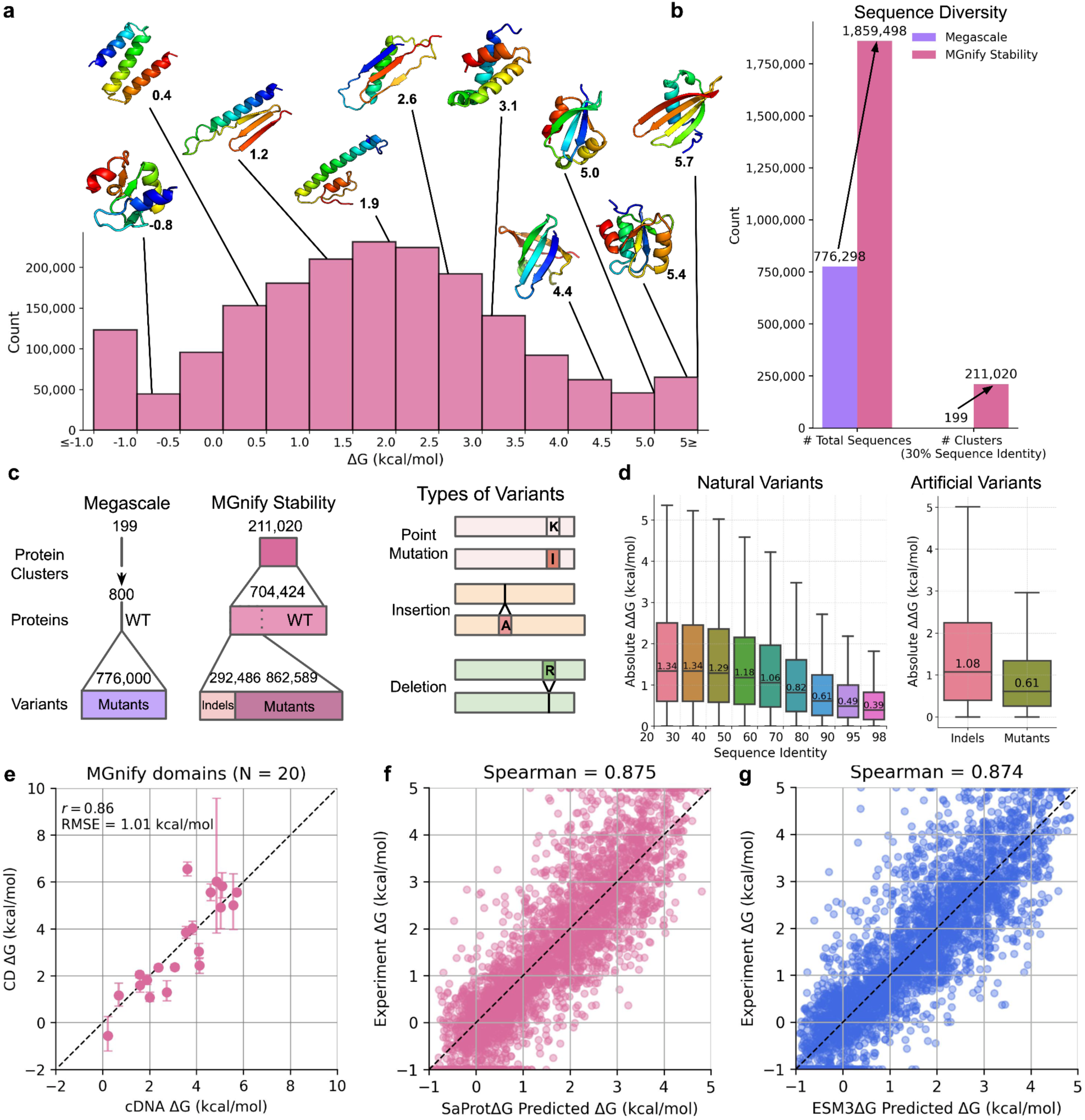
The MGnify Stability dataset contains >1.8 million sequences and >200,000 unique protein clusters with experimentally measured folding stability. **a,** Distribution of folding stability (ΔG) values for MGnify proteins. Sequences range from 60–80 residues with ΔG spanning -1 to 5 kcal/mol, where higher ΔG indicates greater folding stability. **b,** Comparison of dataset scale between Megascale and MGnify Stability. Clustering at 30% sequence identity yields 199 clusters in Megascale and 211,020 clusters in MGnify Stability, which includes wild-type proteins, point mutants, and indels. **c,** Composition of the MGnify Stability and Megascale datasets. The left panel shows the proportions of wild-type, mutant, and indel sequences. The right panel illustrates the three variant types represented in MGnify Stability: point mutations, insertions, and deletions. **d,** Differences in folding stability within and between sequence groups. The left panel shows median ΔG differences among natural variants within clusters across sequence identity bins (20–98%). The right panel shows median ΔG differences between artificial variants and their wild-type counterparts, with indels in pink and point mutants in green. **e,** Correlation between stability measured by circular dichroism (CD) and the cDNA display method for 20 selected domains from the MGnify Stability dataset. In this plot, N represents the number of domains, and *r* represents the Pearson correlation coefficient. Representative CD analysis is shown in Figure S2. **f,** Performance of the SaProt sigmoid LoRA (SaProtΔG) and **g,** ESM3 sigmoid LoRA (ESM3ΔG) models on the MGnify Stability test dataset of 3,283 sequences. The scatter plot compares predicted versus experimental ΔG values obtained from cDNA display proteolysis.

### Measuring stability for diverse domains at scale

To expand the scale and diversity of experimental folding stability measurements, we analyzed a large and diverse collection of domains from the MGnify metagenomic database (Richardson et al. 2023). Because full proteins often contain multiple domains that fold independently (each with their own stability), putative domains 60-80 amino acids in length were extracted from full-length MGnify proteins by clustering the predicted aligned error (PAE) matrices from AlphaFold2 predicted structures. Domains were filtered on several criteria, including AlphaFold2 pLDDT confidence, sequence length, predicted secondary structure, and hydrophobicity. We further expanded this set by generating 710,605 single-substitution variants, 150,095 double-substitution variants, 1,889 variants with more than two substitutions, and 292,486 insertion and deletion (indel) variants of these sequences (Fig. 1c). These domains are highly diverse: clustering the library with MMseqs2 at 30% identity yields 211,020 clusters, and clustering with a TM-score threshold of 0.5 yields 129,303 structural clusters. This represents a vast expansion in sequence and structural diversity relative to the previous Mega-scale folding stability dataset (Fig. 1a,b)

All sequences were encoded in three oligo pools containing 573,641–899,998 sequences and assayed for folding stability using our previously developed cDNA display proteolysis technique (Tsuboyama et al. 2023), creating the MGnify Stability Dataset. In cDNA display proteolysis, all proteins are expressed together in one mixture using cell-free protein synthesis and cDNA display, then proteins are challenged with different concentrations of proteases. Protease cleavage rates are inferred by sequencing the recovered cDNA from intact (uncleaved) protein-cDNA complexes, and folding stabilities are computed by comparing each sequence’s experimental cleavage rate to its predicted cleavage rate in its unfolded state based on a calibration model (Tsuboyama et al. 2023).

Our previous work extensively examined the consistency between cDNA display proteolysis and traditional folding stability experiments using purified proteins, finding correlations (Pearson’s *r*) within groups of related mutants ranging from 0.75 to 0.96 (Tsuboyama et al. 2023). Here, we further validated these measurements in several ways. As before, all sequences were analyzed in two separate, independent experiments: once using the protease trypsin and once using the orthogonal protease chymotrypsin. Folding stabilities were highly consistent between these independent experiments (RMSE 0.88 kcal/mol, 0.70 kcal/mol after removing a global offset, Pearson’s/Spearman’s correlations 0.89/0.90, Supplementary Fig. 1). Replicates of 1,997 sequences in multiple libraries were also highly consistent after a calibration procedure (RMSE 0.36 kcal/mol). Sequences with higher sequence identity tended to have more similar folding stabilities (median difference 0.4 kcal/mol for 95-98% identity, versus 1.3 kcal/mol below 30% identity, Fig. 1d). Finally, we individually purified 20 domains and measured their folding stabilities by GdnHCl chemical denaturation and the linear extrapolation method (Pace and Alexander 1971) (Supplementary Fig. 2). Overall, these stabilities were very consistent with our high-throughput results (RMSE 1.0 kcal/mol, Pearson *r* 0.86, removing one outlier improves RMSE to 0.79 kcal/mol and Pearson *r* to 0.91, Fig. 1e, Supplementary Fig. 2)

### Predicting stability using existing models

How well can simple biophysical properties and existing models predict our stability measurements? We used a subset of the MGnify Stability Dataset (our testing set of 3,283 sequences from 590 sequence clusters) to analyze correlations between experimental stability and a range of biophysical features and “zero-shot” outputs from existing machine learning models. We computed biophysical properties of these sequences using a previously developed featurization pipeline (Kim et al. 2022; Ferrari et al. 2025). Out of 2,065 different features, the strongest Spearman correlation was with the amount of buried nonpolar surface area (NPSA) from hydrophobic residues (Spearman’s correlation 0.60), which was stronger than the correlation from total buried NPSA (0.56) or the count of nonpolar residues (0.48) (Fig. 2a). This result reinforces the “dominant” role of buried hydrophobic interacts in folding stability (Dill 1990a). Still, this correlation is far from perfect, as illustrated by the many domains with minimal buried NPSA yet high folding stability, as well as the reverse (Fig. 2b).

**Figure 2.**
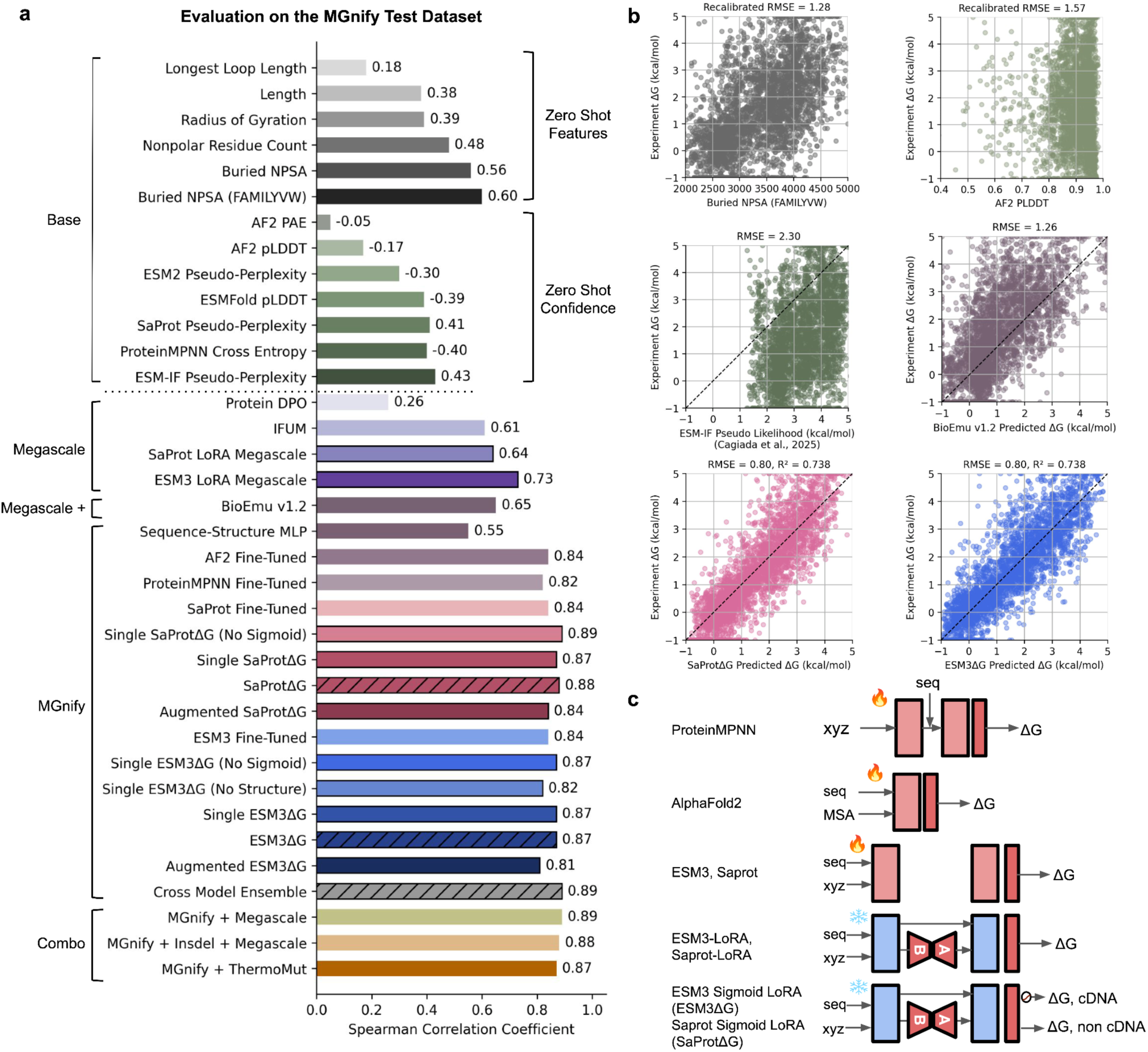
Structure-informed language models achieve superior absolute stability prediction performance. **a,** Spearman correlations between predicted and experimental ΔG values for different model categories. Models are grouped into five classes: zero-shot protein features, zero-shot prediction confidence scores, models trained solely on the Megascale dataset, models trained solely on the MGnify Stability dataset, and SaProt LoRA models trained on combined datasets (MGnify Stability, Megascale, MGnify Stability indels, and ThermoMut). Hatch marks indicate ensemble models obtained by ensembling three independently trained models. **b,** Performance on the MGnify Stability test dataset for representative models. Scatter plots compare experimental ΔG values (y-axis) with predicted ΔG values (x-axis) for six methods: Buried NSPA(FAMILYVW), AF2 pLDDT, ESM-IF pseudo-likelihood (Cagiada et al., 2025), BioEmu v1.2, SaProtΔG, and ESM3ΔG trained on the MGnify Stability dataset. **c,** Architectures of fine-tuned protein language models. ProteinMPNN, AlphaFold2, ESM-3, and SaProt were fine-tuned with additional per-residue stability prediction heads and confidence heads. For full fine-tuning, model weights were unfrozen and trained alongside these heads. For LoRA-based training, ESM-3 and SaProt were trained with LoRA adapters and stability prediction heads, where only the LoRA parameters and heads were trainable, while original model weights remained frozen. Pink modules (indicated by the fire symbol in the figure) represent trainable weights that were updated during training, whereas blue modules (indicated by the ice symbol) represent frozen weights that were not updated.

Structure-based sequence likelihood methods such as ProteinMPNN Cross-Entropy (Dauparas et al. 2022) and ESM-IF pseudo-perplexity (Hsu, Verkuil, et al. 2022) showed weaker correlations with stability than simply calculating the total buried NPSA from hydrophobic residues on the predicted structure (Spearman correlation of 0.43 for ESM-IF pseudo-perplexity, the best zero-shot baseline model, Fig. 2a). Likewise, AlphaFold2 and ESMFold pLDDT showed minimal correlations with experimental stability (Spearman correlations of -0.17 and -0.39, respectively, Fig. 2a), as expected because these domains were prefiltered for high pLDDT. These low correlations highlight the difference between a sequence’s likelihood of occurring naturally (influenced by evolutionary processes, including selection for function and solubility) and its thermodynamic folding stability (Tsuboyama et al. 2023). Overall, existing methods showed limited accuracy at predicting stability without specific training on large-scale experimental stability measurements.

### Fine-tuning structure-informed models improves absolute stability prediction

To leverage our large-scale experimental data for predicting absolute folding stability, we fine-tuned deep learning models pre-trained for sequence and structure prediction tasks. Pre-training on massive protein databases captures general biophysical regularities, enabling efficient adaptation to downstream biophysical objectives such as stability prediction. Our training set for fine-tuning consisted of 960,216 MGnify stability sequences with 525,127 wild-type and 435,089 point mutants (indel mutations were not used for training). All training sequences had under 30% sequence identity to our testing set used above (the same test set is used throughout Fig. 2A). We performed fine-tuning by adding a regression head that aggregated per-residue stability estimates into a global ΔG prediction (Fig. 2c). To further improve accuracy, we jointly trained models on absolute stabilities (ΔG) and relative stability changes (ΔΔG) from mutations, combining losses from wild-type ΔG, mutant ΔG, and ΔΔG predictions with respective weights of 0.3, 0.3, and 1.0 (Methods). This dual training scheme enabled the model to integrate local mutational constraints with global determinants of folding stability. Among several fine-tuning approaches tested (Fig. 2c), parameter-efficient fine-tuning with low-rank adaptation (LoRA) achieved the best performance (Fig. 2a, middle, dark pink box). The resulting models, SaProtΔG and ESM3ΔG, achieved Spearman correlations of 0.88 and 0.87, respectively, with RMSE values of 0.80 kcal mol⁻¹ for both, establishing a new benchmark for absolute stability prediction on the MGnify Stability test set (Fig. 2a,b). Both models are ensembles of three independent training runs; results from one training run are denoted “single” in Fig. 2a. Notably, training on the MGnify Stability dataset yielded substantially higher predictive accuracy than training on the previous Megascale (mutational) dataset alone, underscoring the value of large, diverse absolute stability measurements for model generalization (Fig. 2a,c).

In addition to the “base” SaProtΔG and ESM3ΔG models, we also developed “augmented” versions of these models aimed at minimizing potential bias from training on cDNA display proteolysis data. In the MGnify Stability dataset, domains with less secondary structure near the termini showed slightly decreased average folding stability (0.1-0.3 kcal/mol), potentially due to low level cleavage from the folded state (Supplementary Fig. 3). This bias in the training data may lead SaProtΔG and ESM3ΔG to underestimate folding stability for protein structures with loop segments at the termini. The augmented models mitigate this bias by (1) training on a more limited subset of data, and (2) augmenting this data with artificially added terminal segments (Methods, Supplementary Fig. 3). However, these augmented models have weaker performance on the MGnify Stability test set, because any bias in the proteolysis measurements affects the testing data as well as the training data (Fig. 2a, Supplementary Fig. 4). As described further on, the Augmented ESM3ΔG model improved performance at predicting stability measured by traditional methods (not proteolysis).

Due to the strong correlation between experimental stability and buried NPSA from hydrophobic amino acids, we examined whether SaProtΔG, ESM3ΔG, and other models retained predictive accuracy on a subset of the testing set with high buried NPSA. Restricting the test set to domains with buried NPSA > 3,000 Å^2^ (2,006 total domains, 61 % of the testing set) reduces the Spearman correlation between buried NPSA from nonpolar amino acids and experimental stability from 0.60 to 0.36, but the SaProtΔG and ESM3ΔG correlations remain strong (from 0.88 to 0.80 and 0.88 to 0.77, respectively, Supplementary Fig. 5). Sequence-likelihood models such as SaProt pseudo-perplexity and ESM-IF pseudo-perplexity lose almost all correlation with stability on this high buried NPSA subset (Supplementary Fig. 5). To further assess whether performance depends on similarity to training data, we divided the test set into domains with high structural similarity to the training set (TM-score > 0.5) and low similarity (TM-score < 0.5) and compared their errors across the ΔG range. Absolute prediction errors are broadly comparable, with modestly lower errors for TM-score > 0.5, indicating that performance largely generalizes beyond close structural homologs (Supplementary Fig. 6).

### Accurate prediction across sequence variation

Most protein stability modeling focuses on predicting the effects of single amino acid substitutions (ΔΔG) based on the local structural environment of the mutation. These approaches assume that stability changes are dominated by local interactions, and therefore struggle to capture substitutions that induce larger conformational rearrangements, multiple simultaneous mutations, or insertions and deletions. Predicting mutational effects directly from absolute stability estimates (Fig. 3a) offers a more general framework, as it allows conformations to be evaluated independently and places no restrictions on the number or type of sequence variations considered. However, this approach remains challenging because errors in absolute stability predictions may not cancel between wild-type and mutant sequences, potentially amplifying ΔΔG uncertainty.

**Figure 3.**
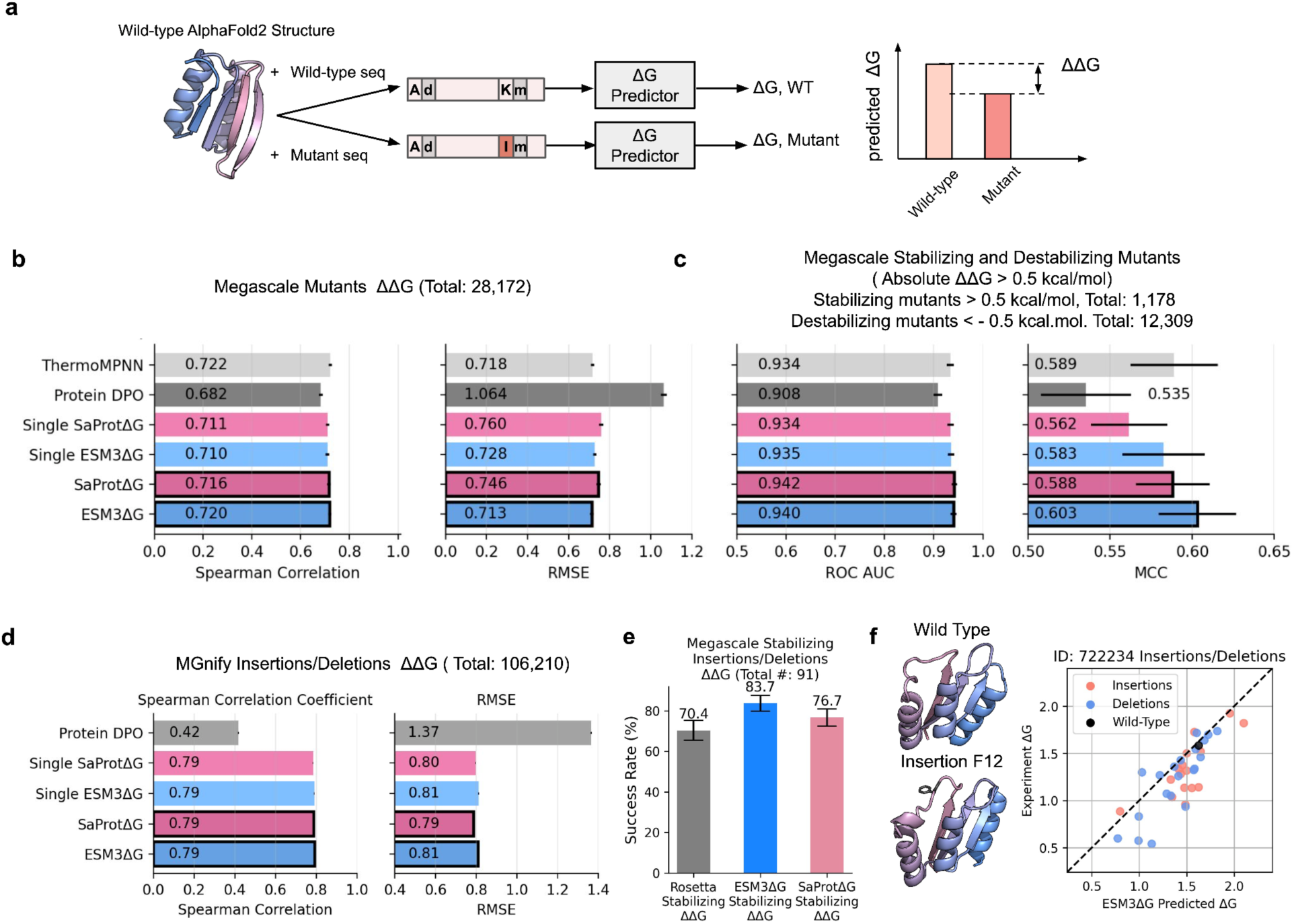
Fine-tuned stability language models accurately predict the effects of point mutations and insertions/deletions. **a,** Schematic of ΔΔG prediction. For a given backbone structure, wild-type and mutant sequences are evaluated independently to predict ΔG, and ΔΔG is calculated as the difference between mutant and wild-type values. **b,** Model performance on Megascale point mutants. Predictions for 28,172 mutants were evaluated by Spearman correlation and RMSE. Error bars indicate uncertainty across bootstrap resamples. **c,** Model predictions on the Megascale stabilizing/destabilizing dataset (12,309 destabilizing mutants and 1,178 stabilizing mutants) were evaluated using AUC and MCC. Variants were defined as stabilizing if ΔΔG > 0.5 kcal/mol to reduce noise. Error bars indicate uncertainty across bootstrap resamples. **d,** Model ΔΔG predictions on the MGnify insertion and deletion dataset (Total: 106,210 variants). Error bars indicate uncertainty across bootstrap resamples**. e,** Success rates for predicting stabilizing insertions and deletions. Performance of SaProtΔG, ESM3ΔG was compared with Rosetta physics-based evaluation on 91 sequences from the Megascale stabilizing indel dataset. Error bars represent bootstrap estimates over 1,000 resamplings. **f,** The left panel shows a stabilizing insertion of phenylalanine at position 12 (ID: rocklin_batch2_722234). Phenylalanine at position 12 is colored black. The right panel shows a scatter plot of experimental versus ESM3ΔG-predicted ΔG values. The corresponding wild-type ΔG was 1.62 kcal/mol. Blue points represent deletions; pink points represent insertions.

To test whether absolute stability predictors can accurately capture the impact of substitutions, insertions, and deletions on folding stability (ΔΔG), we evaluated both SaProtΔG and ESM3ΔG at predicting the effects of point mutants measured in the original Megascale dataset. The ThermoMPNN model (a model trained to predict ΔΔG using the first Megascale experimentally dataset and features extracted from ProteinMPNN, (Dieckhaus et al. 2024; Dauparas et al. 2022) previously introduced a testing set of 28,172 single-point mutants across 28 wild-type proteins, and we re-used this set for our benchmarking (Supplementary Fig. 7). SaProtΔG and ESM3ΔG both models achieved Spearman correlations and RMSE values comparable to ThermoMPNN (Fig. 3b). Both models performed better than ProteinDPO (Widatalla et al. 2024), a structure-conditioned language model based on ESM-IF (Hsu, Verkuil, et al. 2022) and fine-tuned on Megascale data (Fig. 3b). SaProtΔG and ESM3ΔG also showed comparable sensitivity in identifying stabilizing mutations, reaching AUCs (Areas Under the Curve) of 0.94 (compared to 0.93 for ThermoMPNN) and Matthews correlation coefficients (MCC) of 0.58 (SaProtΔG) and 0.60 (ESM3ΔG), compared to 0.59 for ThermoMPNN (Fig. 3c). Although 9 of the 28 wild-type proteins in the ThermoMPNN test set have sequence identity > 0.3 to proteins in the MGnify Stability training set, SaProtΔG and ESM3ΔG performance remains similar on the mutant sets from the remaining 19 wild-type proteins (21,118 single point mutants, Supplementary Fig. 8).

SaProtΔG and ESM3ΔG also performed well at predicting effects of insertions and deletions in MGnify domains (Fig. 3d). The RMSEs for predicting indel effects in MGnify domains (0.79 and 0.81 kcal/mol for SaProtΔG and ESM3ΔG) were only marginally higher than the RMSEs for predicting substitution effects (0.76 and 0.71 kcal/mol, (Fig. 3b)). On a benchmark of stabilizing insertions and deletions curated from the Megascale dataset in previous work (Gutierrez and Rocklin 2024), ESM3ΔG reached 83.7% classification accuracy—a 13-percentage-point improvement over the 70.4% accuracy previously reported using the Rosetta energy functions (Fig. 3e). One illustration is shown in Figure 3f, highlighting accurate ESM3ΔG predictions of stabilizing insertions and deletions in a mixed α/β domain from the MGnify Stability set. Substitution-based models like ThermoMPNN are architecturally incapable of predicting these effects. Although ProteinDPO (a structural-likelihood-based method) can evaluate indel variants, the overall correlation with variation in folding stability was low (Spearman 0.42, Fig. 3d). Together, these results demonstrate that absolute stability prediction provides a unified and general framework capable of modeling diverse sequence variations beyond point mutations.

### Limitations of predictors for large or highly stable domains

We next analyzed how well SaProtΔG and ESM3ΔG could generalize to predict ΔG and ΔΔG measurements on larger and more stable proteins than those in our MGnify Stability dataset. Existing ΔΔG benchmarking sets such as S669 (Pancotti et al. 2022; Xavier et al. 2021) suffer from significant data quality problems, including measurements of irreversible unfolding, non-two-state proteins, and ΔΔG of binding measurements mixed in with ΔΔG of folding. We therefore curated our own dataset of 1,724 folding stability measurements on individually purified proteins from ThermoMutDB (Xavier et al. 2021)). This ‘S1724’ includes mutants from 35 wild-type proteins with folding stabilities from 3 to 13 kcal/mol that all unfold reversibly in a two-state manner (see Methods). To accommodate the differences in experimental conditions and imbalances between the number of mutants measured for each wild-type, we evaluated performance separately for each group of mutants from the same wild-type and used bootstrapping to estimate confidence intervals of all quality metrics.

Because these proteins have a wider folding stability range than our MGnify Stability training set, we modified SaProtΔG and ESM3ΔG to extend the range of their stability predictions. Both models (and their augmented versions) use sigmoidal transformation at the prediction head to constrain predictions to the range of -1 to 5 kcal/mol; we disabled this transformation for all four models for evaluating performance on this S1724 set. The Augmented ESM3ΔG model in particular showed improved performance at predicting ΔG and ΔΔG on the new S1724 benchmark, while the Augmented SaProtΔG model showed minimal improvement over the original SaProtΔG model. Because the Augmented ESM3ΔG model had the strongest performance on the larger proteins in S1724, we focused our remaining analysis (Fig. 4-6) on this model.

**Figure 4.**
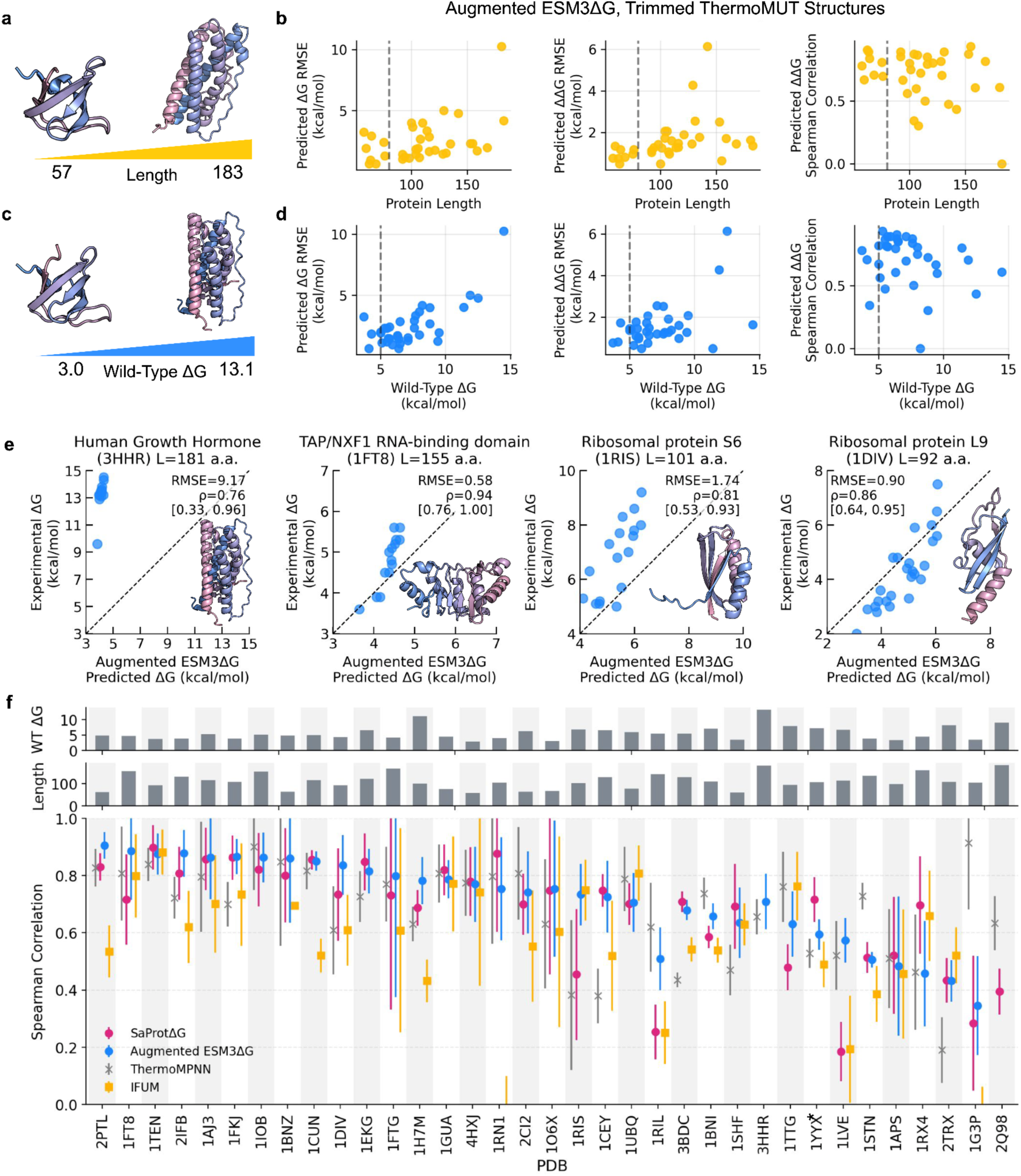
ESM3ΔG achieves comparable performance to ThermoMPNN on independent datasets but shows limitations for highly stable and large proteins. Performance on the ThermoMut dataset. **a,** Schematic showing the protein length range and representative structures. **b,** Augmented ESM3ΔG predictions for each set of mutants from ThermoMut. Each point represents predictions for one wild-type protein and its mutants. Performance metrics (ΔG RMSE, ΔΔG RMSE, and Spearman’s rank correlation) are plotted against increasing protein length (x-axis). A vertical dotted line indicates 80 residues, the maximum protein length in the training data. **c,** Schematic showing the wild-type ΔG range of ThermoMut proteins. **d,** Stability distribution of ThermoMut proteins, with wild-type stabilities spanning 3.0 - 13.1 kcal/mol, extending beyond the training range of ESM3ΔG; model performance metrics are plotted against increasing wild-type ΔG values. A vertical dotted line indicates 5 kcal/mol, the maximum wild-type stability in the training data. **e,** Examples of Augmented ESM3ΔG predicted ΔG versus experimental ΔG for four proteins: 3HHR, 1FT8, 1RIS, and 1DIV. The RMSE, Spearman correlation, and the lower and upper bounds of the confidence interval for the Spearman correlation are shown on the plot. **f,** Spearman’s rank correlation between predicted and experimental stability for individual wild-type PDB structures. Correlations are shown for predictions from SaProtΔG, ESM3ΔG, and IFUM. Each correlation was bootstrapped 1,000 times, with error bars indicating the standard deviation across replicates; protein length and wild-type ΔG are shown above each data point. * on 1YYX indicates high sequence identity with the MGnify Stability training dataset.

Augmented ESM3ΔG performance varied from protein to protein, with ΔG and ΔΔG accuracy lower for the groups of mutants from the largest and most stable wild-type domains (Fig. 4a-d). The accuracy of ΔG prediction was especially low for groups of mutants with wild-type stability above 10.0 kcal/mol, highlighting the need for improved training data on these high-stability proteins. This limitation is exemplified by human growth hormone (3HHR, wild-type ΔG = 13.1 kcal/mol), for which predicted and experimental ΔG values differed substantially (Fig. 4e). Although the training data had a maximum domain length of 80 amino acids, Augmented ESM3ΔG achieved reasonably accurate ΔG and ΔΔG predictions for many longer domains (ΔΔG RMSE < 2 kcal/mol and Spearman’s rank correlation coefficient > 0.7); examples ranging from 92 to 155 amino acids in length are shown in Fig. 4e. Removing the sigmoid transformation from the final prediction head expanded the dynamic range of predicted stabilities, improving predictions for proteins such as the Len light chain variable domain (1LVE, 7.7 kcal/mol, (Raffen et al. 1999)), and apocytochrome b562 (1YYX, 9.5 kcal/mol (Feng et al. 2004)), Supplementary Fig. 9). Augmented ESM3ΔG performed similarly to ThermoMPNN at ranking ΔΔG effects across our evaluation set (Fig. 4f). However, these comparisons are imprecise owing to the low number of literature stability measurements and the (often) low dynamic range of ΔΔG measurements, as reflected by the wide 95% bootstrapped confidence intervals in Fig. 4f.

### Conformational sensitivity of stability predictions

By definition, two-state models of folding stability group together all folded conformations (even very different ones) into a single “folded state”. In cDNA proteolysis experiments, multiple different conformations can all contribute to a domain’s measured folding stability, so long as those conformations are substantially more protease resistant than the unfolded state. Still, because SaProtΔG and ESM3ΔG are trained using a single input structure for each sequence, the predictions may be sensitive to the input structure, and the models may even learn to discriminate between higher and lower stability conformations of the same protein, as shown previously (Cagiada et al. 2025). To examine this, we tested the Augmented ESM3ΔG model on two examples of proteins with multiple conformational states taken from Cagiada et al. 2025 (Supplementary Fig. 10). In T4 lysozyme, NMR relaxation dispersion experiments identified an excited state that is less stable than the ground state by approximately 2.0 kcal mol⁻¹ (Bouvignies et al. 2011; Mulder et al. 2001). Augmented ESM3ΔG predicts a stability difference of 1.2 kcal mol⁻¹ between the E and G states, correctly recovering the relative ordering of conformational stabilities. Similarly, for cyclophilin A, NMR relaxation dispersion experiments identified an excited conformational state with a free energy difference of ∼1.7 kcal mol⁻¹ between the two states (Eisenmesser et al. 2005). Augmented ESM3ΔG predicts a ΔΔG of 1.0 kcal mol⁻¹. Although the predicted magnitudes are smaller than the experimental values by 0.5-0.8 kcal mol⁻¹, the model correctly captures the relative stability relationships between conformations given only the PDB structures and identical sequences, highlighting its potential utility for discriminating between alternative conformational states.

### Predicted folding stability correlates with organismal optimal growth temperature

Due to the limited scale and diversity of published folding stability measurements like those in Fig. 4, we sought a larger-scale method for benchmarking our model. We reasoned that organismal optimal growth temperature (Martin K. M. Engqvist 2018) could be an accessible, large-scale proxy measure for folding stability, especially when comparing sequences within homologous families of proteins. It is well established that proteins from thermophilic and hyperthermophilic organisms typically have increased folding stability at room temperature (Luke et al. 2007; Rees and Robertson 2001; Szilágyi and Závodszky 2000; Petsko 2001).

To evaluate how Augmented ESM3ΔG-predicted stabilities correlate with optimal growth temperature, we compiled a test dataset of three million sequences from 583 CATH (structural) families in TED (The Encyclopedia of Domains) (Lau et al. 2024). Within each CATH family, we selected at least 1,000 sequences from organisms spanning a range of optimal growth temperatures (Martin Karl Magnus Engqvist 2018), oversampling thermophilic and hyperthermophilic organisms to improve the uniformity of the growth temperature distribution (Fig. 5a,b, Methods). These domains represented diverse fold classes (Fig 5c), with average pairwise sequence identities within each family ranging from 50-60%. On average, Augmented ESM3ΔG predicted higher folding stabilities for domains from organisms with higher optimal growth temperatures (OGT) for nearly all protein families (Fig. 5d,e,f). Positive slopes (ΔG increment per temperature range) were observed in both small and large domains (Fig. 5f), demonstrating that Augmented ESM3ΔG generalizes beyond its training data of domains <80 residues to capture thermophilic stability trends in larger proteins in a zero-shot manner (i.e. without training to predict optimal growth temperature).

**Figure 5.**
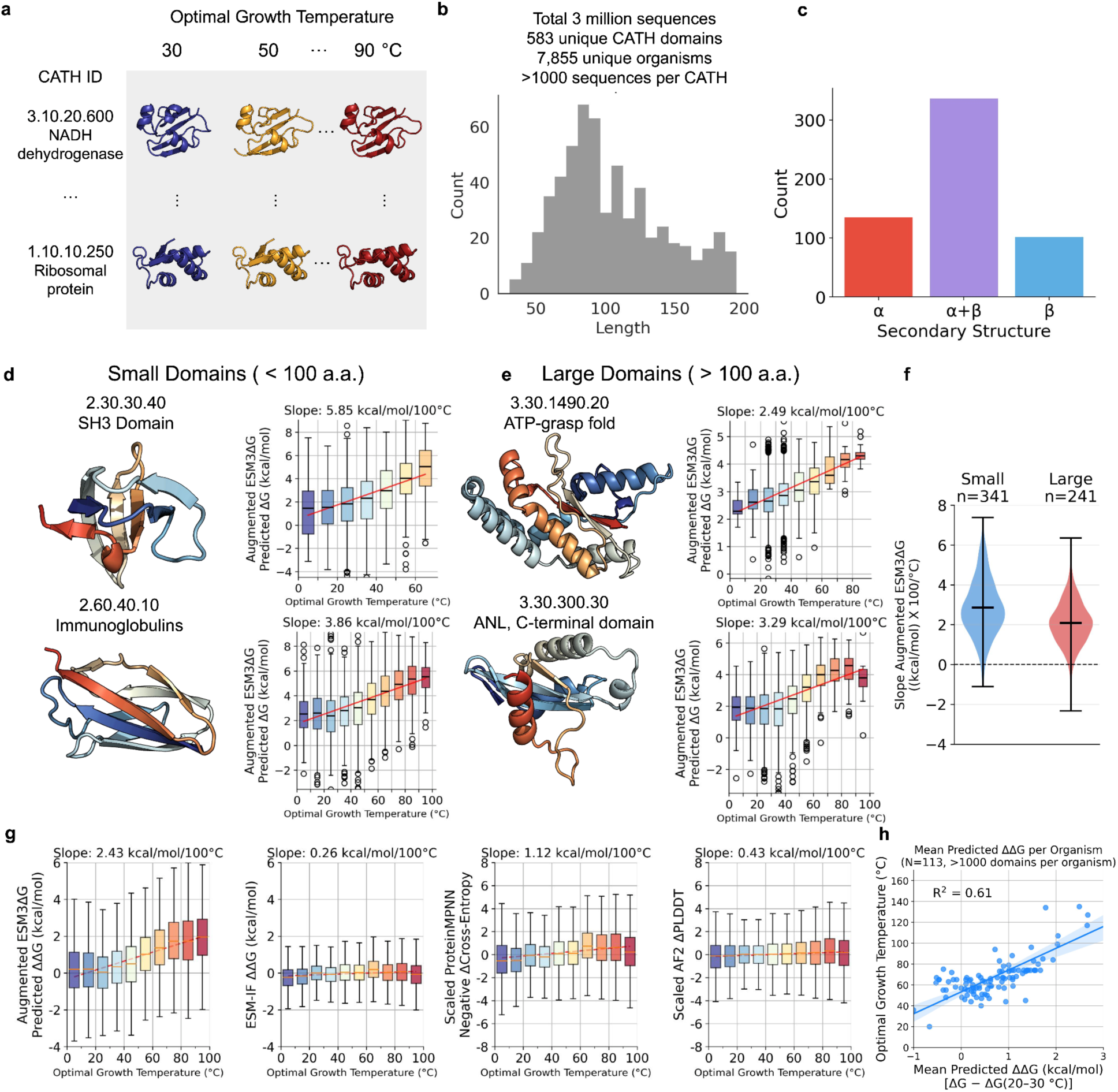
ESM3ΔG can estimate optimal growth temperature using The Encyclopedia of Domains dataset. **a,** Schematic of the TED dataset. Proteins with the same CATH ID but originating from organisms with different OGTs were collected. **b,** Filtering pipeline for TED domains. The final dataset includes 583 unique CATH domains, 7,855 organisms, and >1,000 sequences per CATH domain, restricted to proteins ≤200 residues. **c,** Structural distribution of the dataset. Counts of α, α/β, and β domains are shown. **d,** Stability predictions for small domains. Box plots of ESM3ΔG-predicted ΔG values (y-axis) binned by OGT (10 °C intervals, x-axis) are shown for SH3 domains (CATH 2.30.30.40, top) and Immunoglobulin-like domains (CATH 2.60.40.10, bottom). **e,** Stability predictions for large domains. Distributions are shown for the ATP-grasp fold (CATH 3.30.1490.20, top) and the ANL C-terminal domain (CATH 3.30.300.30, bottom). Titles report slopes of predicted ΔG versus OGT (units: kcal/mol/100 °C). **f,** Distribution of slopes across domain size. Box plots compare slopes of predicted ΔG versus OGT for small (<100 residues) and large (100–200 residues) domains. **g,** Comparison of predictors across all domains. ΔΔG values were calculated as the difference between each variant and the mean stability of mesophilic sequences (OGT = 20–30 °C). Box plots show relationships between OGT and ESM3ΔG ΔΔG, ESM-IF ΔΔG, scaled ProteinMPNN negative cross-entropy, and scaled AlphaFold2 pLDDT. CE and pLDDT scores were recalibrated to the predicted ΔG scale using quantile-based distribution matching. **h,** Organism-level prediction. Mean predicted ΔΔG per organism (relative to the 20–30 °C mesophilic baseline) plotted against OGT, yielding an R² of 0.61.

After observing a significant correlation between our predicted folding stabilities and optimal growth temperature (OGT), we asked how this correlation compared to other “zero-shot” model outputs such as ProteinMPNN cross-entropy and AlphaFold2 pLDDT. To analyze this across different CATH families, we used the mesophilic sequences (OGT 20–30 °C) from each family to establish baseline average mesophilic predictions (predicted ΔG, predicted cross-entropy, etc) for that family. We then compared predictions for all non-mesophilic sequences against the baseline average mesophilic prediction for their CATH family. After averaging data across all CATH families, we found that Augmented ESM3ΔG predicted a linear slope of 2.43 kcal/mol per 100 °C, approximately 2–9 fold steeper than ESM-IF (Cagiada et al. 2025), ProteinMPNN cross-entropy, and AlphaFold2 pLDDT (Fig. 5g; ProteinMPNN scores and pLDDT scores were rescaled using quantile-based distribution matching to the Augmented ESM3ΔG predictions). This indicates that stability predictions from the Augmented ESM3ΔG model more strongly discriminate growth temperature differences compared to other zero-shot confidence scores commonly used to approximate stability. At the organism level, we also observed a strong correlation between each organism’s OGT and the predicted ΔΔG for sequences in that organism compared to the CATH-specific baseline average mesophilic stability (R² = 0.65, Fig. 5h). This indicates Augmented ESM3ΔG captures an organism-wide signature of thermal adaptation without any training on growth temperature data. Of course, training directly on labeled growth temperature data would likely lead to improved predictions at this benchmark (Jiang et al. 2024). Still, different domains from thermophilic organisms can differ widely in stability, and predicting optimal growth temperature is not sufficient to predict stability, though it can be useful in protein engineering (Jiang et al. 2024).

### Stability filtering improves protein design

Last, we evaluated whether the ESM3ΔG models could improve practical protein design workflows. We focused on three different design challenges: (1) triaging de novo designed proteins for stable folding, (2) ranking nanobodies by apparent thermal stability, and (3) triaging de novo designed binders. To evaluate the model at triaging de novo designed proteins, we analyzed two large datasets of experimentally characterized designs: first, a set of 634 Rosetta designs from 12 different studies published between 2012 and 2021 and previously analyzed for folding by circular dichroism (Garcia et al. 2025, 2026), and second, 20,572 small de novo designed protein domains sampled from different design tools and previously analyzed for folding stability by cDNA display proteolysis (Cho et al. 2025). On the set of Rosetta designs, ESM3ΔG showed a strong ability to discriminate experimentally stable from unstable designs (AUC 0.76 ± 0.02, median and 95 c.i. from bootstrapping), exceeding discrimination by AlphaFold2 pLDDT (0.65 ± 0.02) or ProteinMPNN cross-entropy (0.68 ± 0.02, Fig. 6a,b). This is notable because these Rosetta designs were all experimentally tested as purified proteins (not by cDNA display proteolysis), and these designs are also larger and effectively out of sample for the ESM3ΔG training data (mean domain size 100 amino acids ± 28 std. compared to training on domains < 80 a.a.). Furthermore, these designs all predate AlphaFold2 and ProteinMPNN, eliminating any bias due to pre-filtering using these tools, and were not in those tools’ training sets. On the more recent set of de novo designs from 20,572 designs from (Cho et al. 2025), ESM3ΔG showed a very strong correlation with the quantitative folding stability measurements (Spearman’s ρ = 0.88, RMSE = 0.98 kcal/mol, Fig. 6c,d). Although this could be expected because both the training data and this benchmarking data were collected by cDNA display proteolysis, it remains noteworthy because these de novo designs have minimal sequence identity to any sequences in the training set (7 out of 20,668 sequences at ≥30% identity).

**Figure 6.**
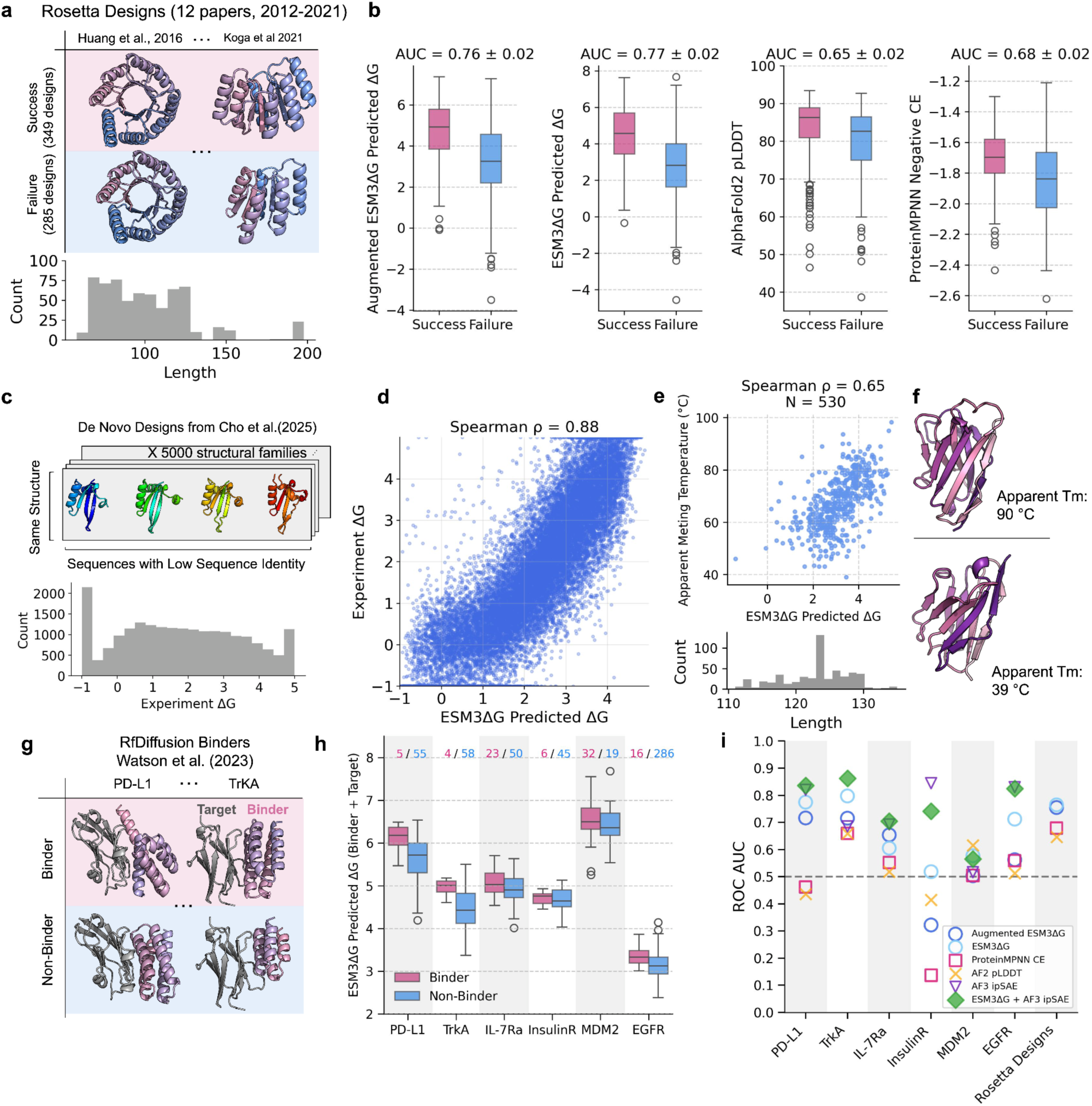
ESM3ΔG predicts the absolute stability of de novo protein designs and serves as an effective score for identifying functional binders. **a,** Rosetta designs collected from 12 published studies (2012–2021) spanning 50–200 residues. Two examples of successful and failed designs are shown from representative papers. **b,** Box plots comparing successful versus failed designs using SaProtΔG-predicted ΔG (left), AlphaFold2 pLDDT scores (center), and ProteinMPNN negative cross-entropy scores (right). Bootstrapped AUC (1,000 resamples) ± error is shown. **c,** De novo designs from Cho et al. (2024), comprising 5,000 structure families with four sequences per family generated using different design models. **d,** Scatter plot of ESM3ΔG-predicted versus experimentally measured ΔG values for Cho et al. designs.**e,** Scatter plot of ESM3ΔG-predicted ΔG versus nanobody apparent melting temperature (Tm). **f,** Nanobodies from NbThermo: a high apparent Tm example (top, 90 °C) and a low apparent Tm example (bottom, 39 °C). **g,** RFDiffusion binders from Watson et al. (2023) across five targets. Representative examples are shown for PD-L1 and TrkA binders (binders in purple; targets in gray). **h,** Box plots of predicted ΔG for binders versus non-binders per target. Analysis includes RFDiffusion targets and EGFR binders from the Adaptyv Bio competition (binders in pink; non-binders in blue). **i,** ROC AUC per target comparing scoring methods: Augmented ESM3ΔG, ESM3ΔG on binder–target complexes, ProteinMPNN cross-entropy (binder only), AlphaFold2 pLDDT (binder only), AlphaFold3 ipSAE minimum score, and a combined score (ESM3ΔG + AlphaFold3 ipSAE minimum), computed by normalizing both scores and summing them.

ESM3ΔG also showed promise at ranking nanobodies by apparent melting temperature. To analyze this, we predicted folding stability for the 530 experimentally characterized nanobodies in the NbThermo dataset (Valdés-Tresanco et al. 2023), a diverse group of sequences originating from 389 unique frameworks. Nanobodies represent a challenging test case for several reasons. First, these single-domain antibodies possess unique structural features including extended CDR3 loops while also having limited evolutionary information compared to natural protein domains (Muyldermans 2013) (Valdés-Tresanco et al. 2023). Second, nanobody melting experiments are often not reversible, and these (equilibrium or not) melting temperatures are not expected to perfectly correlate with equilibrium folding stability at 25°C. Finally, these domains are also larger than the domains in the training data (123 ± 5 a.a., mean ± std.). Despite these challenges, predicted folding stabilities correlated with apparent melting temperatures (Spearman ρ = 0.65; Fig. 6e,f), indicating that ESM3ΔG captures stability determinants that generalize to engineered antibody scaffolds beyond its metagenomic training data. These results suggest that ESM3ΔG and Augmented ESM3ΔG could be used to filter and rank candidate therapeutics in early-stage development. Furthermore, the ESM3ΔG model likely provides a strong foundational model that could be further tuned using nanobody-specific data to improve performance.

Finally, we analyzed whether ESM3ΔG and Augmented ESM3ΔG could effectively rank de novo designed binders from the original RFdiffusion study targeting five different proteins (Watson et al. 2023), as well as from the AdaptyvBio EGFR design competition (Cotet et al. 2025). Because successful binding depends on both the binder’s ability to fold and its interactions with the target, we ranked designs based on the total predicted ΔG of folding for the binder-target complex, not the predicted binding free energy. Overall, stability predictions on binder-target complexes featuring successful designs were systematically higher than predictions for binder-target complexes with inactive designs. ESM3ΔG achieved higher AUCs in distinguishing successful binders from non-binders than AF2 pLDDT and ProteinMPNN cross-entropy across individual target–binder pairs, except in the case of the insulin receptor (Fig. 6g–i). Although this comparison is biased (binders were designed using ProteinMPNN and filtered by AF2 pLDDT prior to experimental testing, limiting the discriminative power of these metrics), the results indicate that ESM3ΔG provides additional discriminatory power. Ranking designs by AlphaFold3 ipSAE generally provided stronger discrimination than ESM3ΔG, except in the case of TrkA.

## Discussion

The original Megascale folding stability dataset and the new MGnify Stability dataset introduced here substantially improve prediction of absolute folding stability for protein domains under 80 amino acids in length (Fig. 2). Despite the physicochemical complexity of folding stability, with sufficient data the SaProtΔG and ESM3ΔG models both achieve mean squared errors below 1 kcal/mol across a broad range of protein domain structures, including for *de novo* designs unlike those used to train the foundational SaProt and ESM3 models (Fig. 6d). Still, we expect these models can likely be further improved through advances in model architectures (Li and Luo 2025), integration of additional data sources, and the adoption of next-generation foundation models such as AlphaFold3, RosettaFold3, or OpenFold3 (Ahdritz et al. 2024; Corley et al. 2025; Abramson et al. 2024).

SaProtΔG and ESM3ΔG represent powerful new tools for protein engineering. These methods do not replace other tools (i.e. pLDDT (Jumper et al. 2021), ipSAE (Dunbrack 2025)) for ranking designed proteins, but are valuable complements bringing orthogonal information to aid optimization and ranking of designed proteins. Furthermore, just as our models benefitted from the pre-training process used to build SaProt and ESM3, our models (and dataset) can serve as new foundation models and pre-training tasks for learning more complicated biophysical phenotypes, including aggregation propensity (Martell et al. 2025; Jarzab et al. 2020), response to desiccation (Kc et al. 2026), allostery (Faure et al. 2022), conformational dynamics (Ferrari et al. 2025), and protein function (Meier et al. 2021; Hsu, Nisonoff, et al. 2022). The foundational role of folding stability in all these processes supports the use of folding stability as a valuable pre-training objective. For tasks where SaProtΔG and ESM3ΔG performed well in a zero-shot setting (meaning without task-specific training data, such as predicting nanobody apparent melting temperature, Fig. 6e), it is likely that small amounts of task-specific training data can lead to significant improvements (Widatalla et al. 2024; Dieckhaus et al. 2024; Stocco et al. 2025), leveraging the foundational understanding of stability conferred by training on the MGnify Stability dataset.

Despite this progress, the MGnify Stability dataset and SaProtΔG and ESM3ΔG models have significant limitations. Most notably, all training data were limited to proteins <80 amino acids in length, and stabilities were only resolved over the approximate range of -1 to 5 kcal/mol. For ease of experimentation, we also excluded proteins with multiple cysteines (including disulfide bonds) from the MGnify Stability dataset. The cDNA display proteolysis method for measuring folding stability also introduces limitations. Importantly, stability will be underestimated for domains that are protease-cleavable in the folded state. Still, this appears to be rare for small domains (Fig. 1e, and (Tsuboyama et al. 2023)), and the smaller training set used to train the Augmented models should mitigate this further. We have also observed that cDNA display proteolysis overestimates folding stability for domains that bind nucleic acids (Ferrari et al. 2025), as expected, and we did not intentionally correct for this effect in training. These data also represent a single set of experimental conditions, and measuring how the chemical environment modulates stability at scale is an important next step.

The limits of the training dataset (particularly in protein length and stability range) constrain the ability of SaProtΔG and ESM3ΔG to generalize, as shown in Fig. 4. Even within the protein length range of our experiments, it is important to note that our training data is designed to broadly sample protein-like structures and sequences, rather than to achieve uniform coverage of possible amino acid sequences. Unusual sequences (e.g. poly-leucine) or structures (e.g. two independent domains connected by a flexible linker, instead of one single domain) are outside the scope of the current models. Collecting new, more broadly representative training data is likely necessary to develop fully general models for predicting folding stability, and improvements to both DNA synthesis technology (for longer domains) and the cDNA proteolysis method itself (overcoming limits of protein size and dynamic range) will accelerate this. Notably, the cost and difficulty of traditional stability measurements on purified proteins also creates a barrier to independently evaluating cDNA display proteolysis-based models; improved methods here would ensure that next generation models minimize overfitting on proteolysis-specific artifacts.

In small scale tests, both SaProtΔG and ESM3ΔG were sensitive to the input structure used for predictions, as shown in Supplementary Fig. 10. This raises theoretical and practical issues. In principle, folding stability is a property of the sequence alone (for a given environment), and folding stability predictions should be independent of any specific input structure. This is because stability measures foldedness in general (here, protease resistance) rather than folding to a specific structure. Still, training our models using AlphaFold2-predicted “native” structures introduced some structure-dependence to the predictions, including some ability to energetically discriminate native and excited conformational states (Supplementary Fig. 10). This suggests it may be possible to exploit folding stability data to develop more accurate models of conformational energetics (Ferrari et al. 2025), especially by using novel architectures that explicitly model full conformational ensembles (Lewis et al. 2025; Lüth et al. 2026; Roney et al. 2025). Practically, our results suggest users of SaProtΔG and ESM3ΔG may want to test predictions using multiple input structures for proteins of interest. Furthermore, SaProtΔG and ESM3ΔG will underestimate folding stability for input structures that would likely be cleaved from the native state (e.g. structures containing long, unstructured terminal segments with trypsin or chymotrypsin cut sites). Underestimating stability (ΔG) may also lead to underestimating mutational effects (ΔΔG), because poorly folded structures can more easily accommodate disruptive mutations. Input structures to SaProtΔG and ESM3ΔG (and even the Augmented models) should be prepared by removing flexible segments from both termini to isolate the fully cooperative, folded structural model.

Our results illustrate that combining large-scale experimental data and modern deep learning models can enable accurate folding stability predictions across a large and diverse space of protein sequences and folds. Can our data and models illuminate protein physical chemistry as well? It is appealing to hunt for unifying principles explaining differences in stability between proteins, but the main principle appears to be the diversity of factors that all contribute significantly to a protein’s net stability and to differences in stability between proteins (Petsko 2001). Quantifying the cumulative effects of diverse forces and thousands of interactions on stability may be more ideally suited for data-driven machine learning than for physics-inspired, additive models (Alford et al. 2017), particularly given the role of entropy and non-additivity in protein biophysics (Dill 1997).

## Acknowledgements

The authors thank Epsilon Molecular Engineering (EME) for providing the cnvK linker for cDNA display, the Rush University Genomics and Microbiome Core Facility for performing DNA sequencing support, and the Northwestern Proteomics Core for mass spectrometry of individually purified proteins. We also thank Michal Zielinski, Rosalia Schneider, Kathryn Tunyasuvunakool, Simon Kohl, and Rob Fergus for providing the sequences in the MGnify Stability dataset as well as for early assistance with machine learning. We further thank Parisa Hosseinzadeh, Steven Lewis, Brian Trippe, Lucas Nivon, James Roney, and many members of the Rocklin and Ovchinnikov labs for helpful discussions and comments on the manuscript, and Állan Ferrari for assistance computing biophysical features of the test set. This work was supported by Northwestern University Startup Funding (G.J.R.), NIH Award R35GM158118 (G.J.R.), the OpenFold consortium (G.J.R.), NIH award T32GM140995 (trainee fellowship to A.O.), JSPS KAKENHI 19J30003 (K.T.), the Human Frontier Science Program Long-Term Fellowship (K.T.), JST PRESTO Grant JPMJPR21E9 (K.T.), National Science Foundation (NSF) NRT Award 2021900 (C.M.P.), and NIH F31GM151811 (C.M.P.). Y.C. acknowledges support from the SBS Scholarship Program and the Takeda Fellowship. S.O. was supported by the National Science Foundation under grant MCB2032259, and Amgen. This research was supported in part through the computational resources and staff contributions provided for the Quest high performance computing facility at Northwestern University which is jointly supported by the Office of the Provost, the Office for Research, and Northwestern University Information Technology.

## Author contributions

Y.C. and K.T. contributed equally to this project. Y.C., K.T., S.O., and G.J.R. conceived the project. K.T. performed and analyzed all cDNA display proteolysis experiments. Y.C. developed the machine learning models and performed all benchmarking and data analysis. T.L. performed and analyzed all circular dichroism experiments, assisted by Q.W. and C.M.P.. M.D.J. expressed all proteins for circular dichroism analysis, with assistance from J.T. and C.M.P.. A.O. compiled the growth temperature benchmarking set. Y.C. and G.J.R. wrote the manuscript with input from all authors. S.O. and G.J.R. acquired funding for the project and supervised the research.

## Data and code availability

The source code and pre-trained model weights are publicly available at GitHub (https://github.com/yehlincho/absolute-stability-predictor) and Hugging Face (https://huggingface.co/Yehlin/absolute-stability), respectively. The full MGnify stability dataset, training dataset, benchmark dataset, and raw data processing scripts are deposited at Zenodo (https://forms.gle/4ZnXZSnTBvaykkAi9).

## Supplementary Figures

**Supplementary Figure 1.**
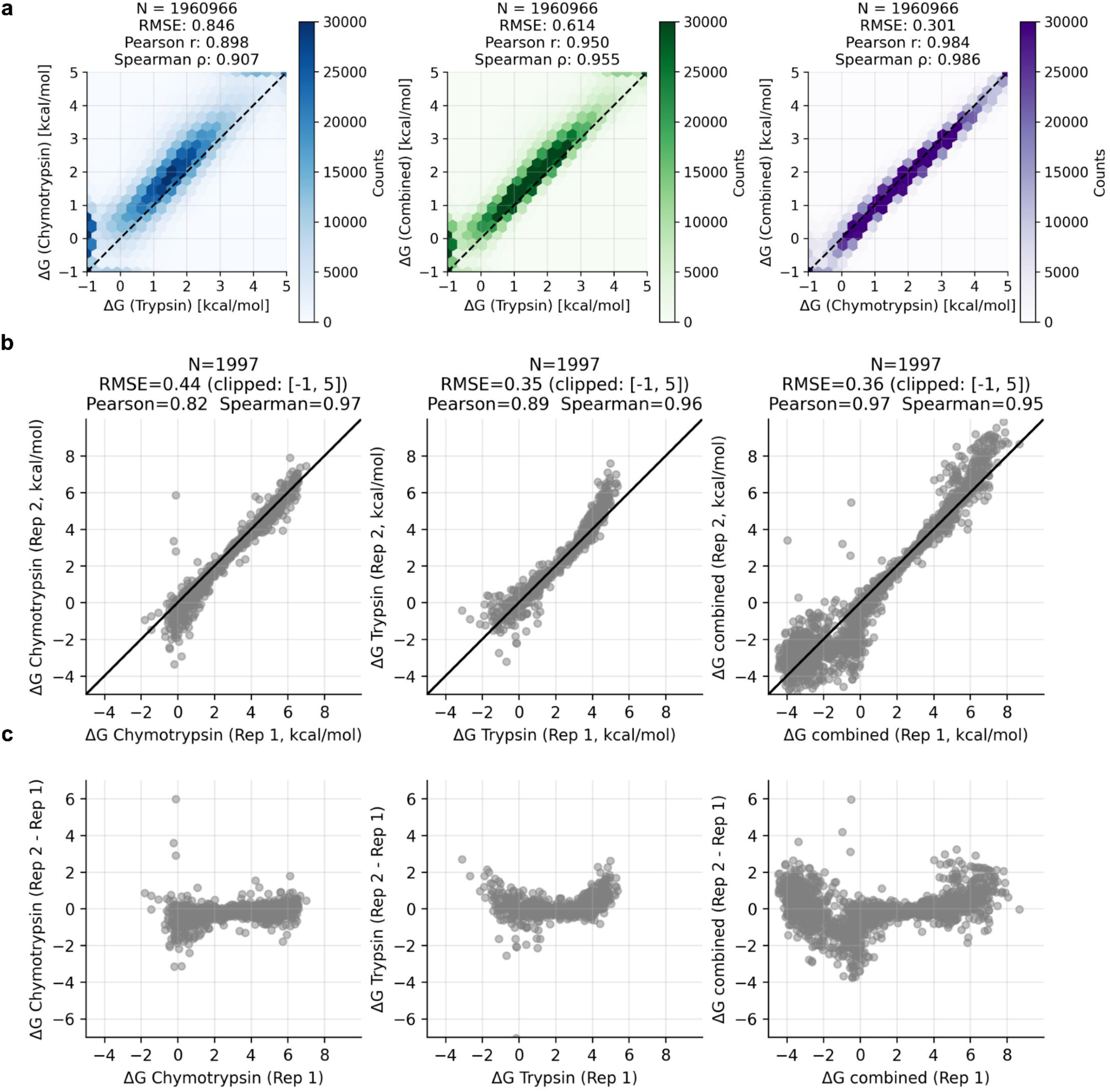
Comparison of ΔG-based stability measurements between trypsin and chymotrypsin and across experimental replicates. **a,** Stability measurements for ∼2 million datapoints obtained using chymotrypsin and trypsin. Left: ΔG (kcal/mol) measured by chymotrypsin versus ΔG measured by trypsin. Middle: combined ΔG values plotted against measurements from trypsin. Right: combined ΔG values plotted against measurements from chymotrypsin. **b,** Reproducibility of stability measurements across a total of 2,003 replicate designs. ΔG values from replicate 2 plotted against replicate 1, with chymotrypsin shown on the left and trypsin shown on the right. **c,** Difference in ΔG between replicate 2 and replicate 1 plotted against replicate 1.

**Supplementary Figure 2.**
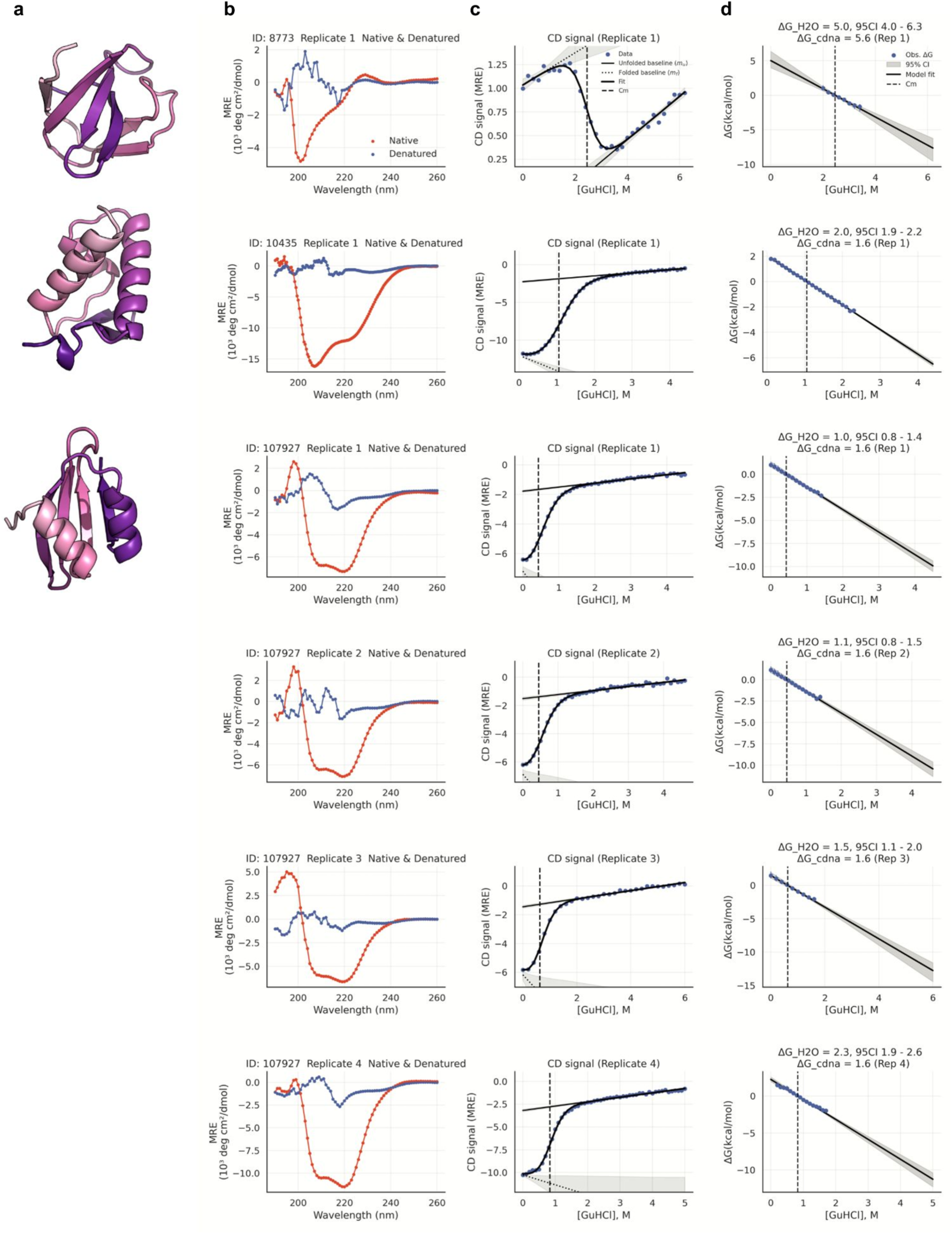
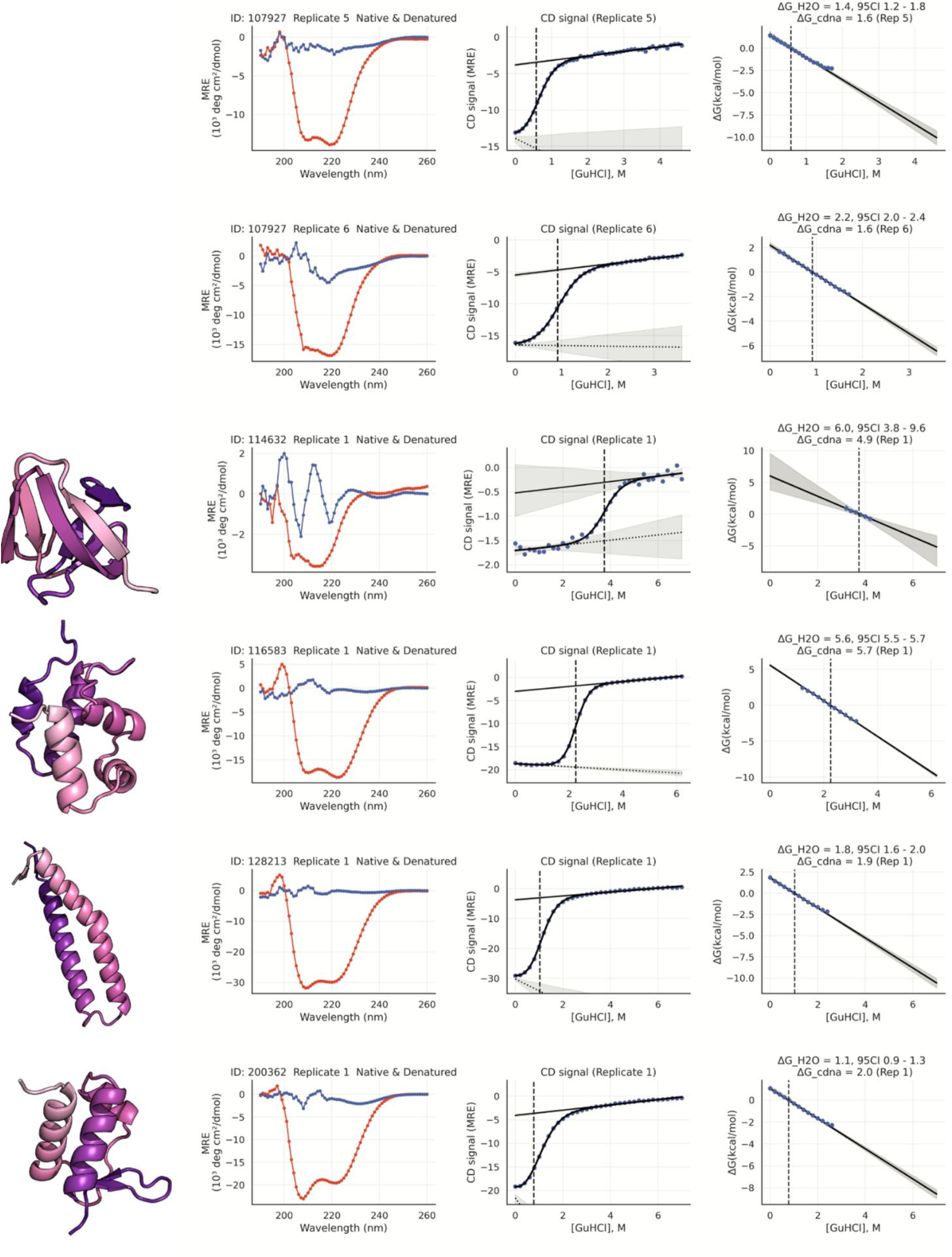
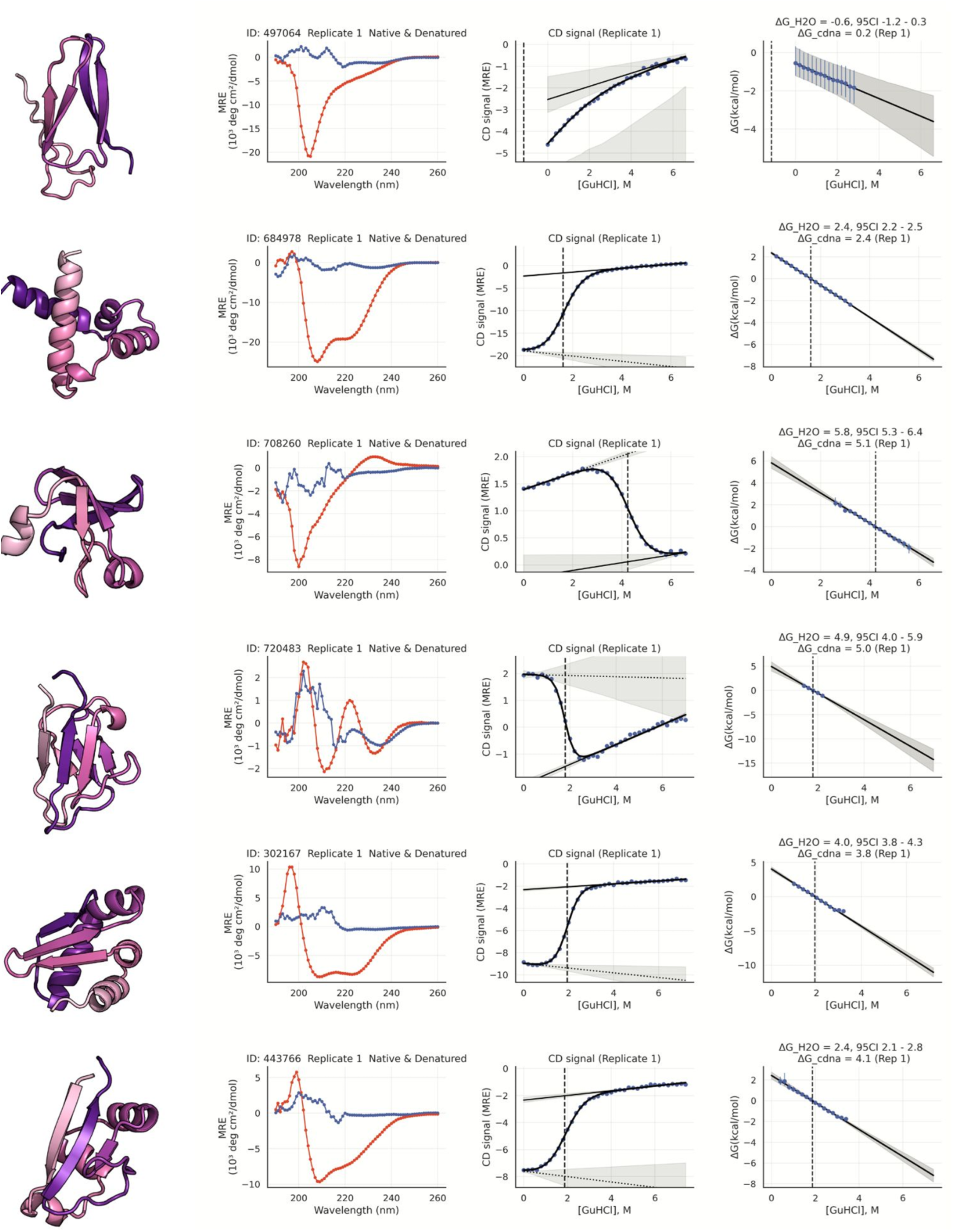
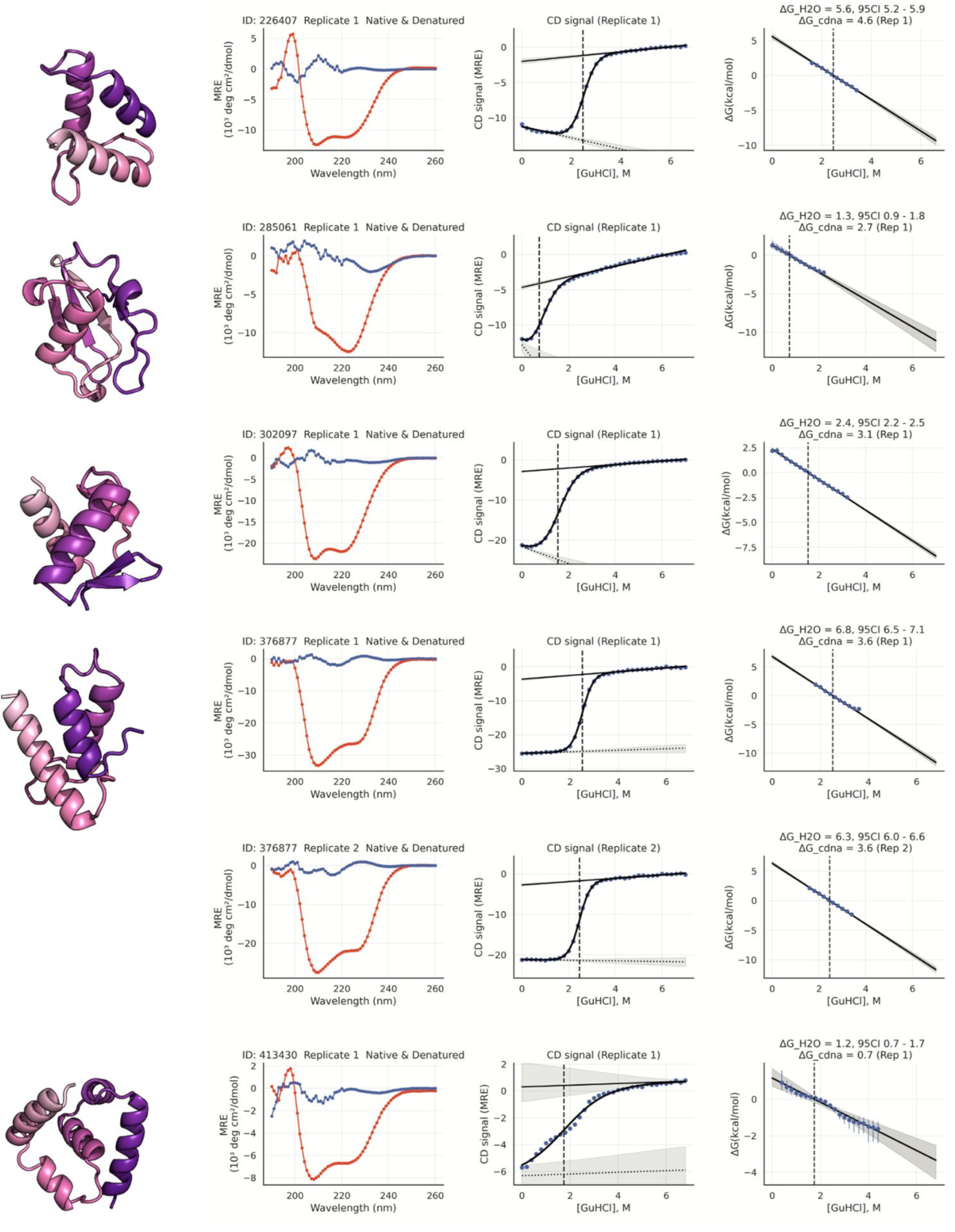
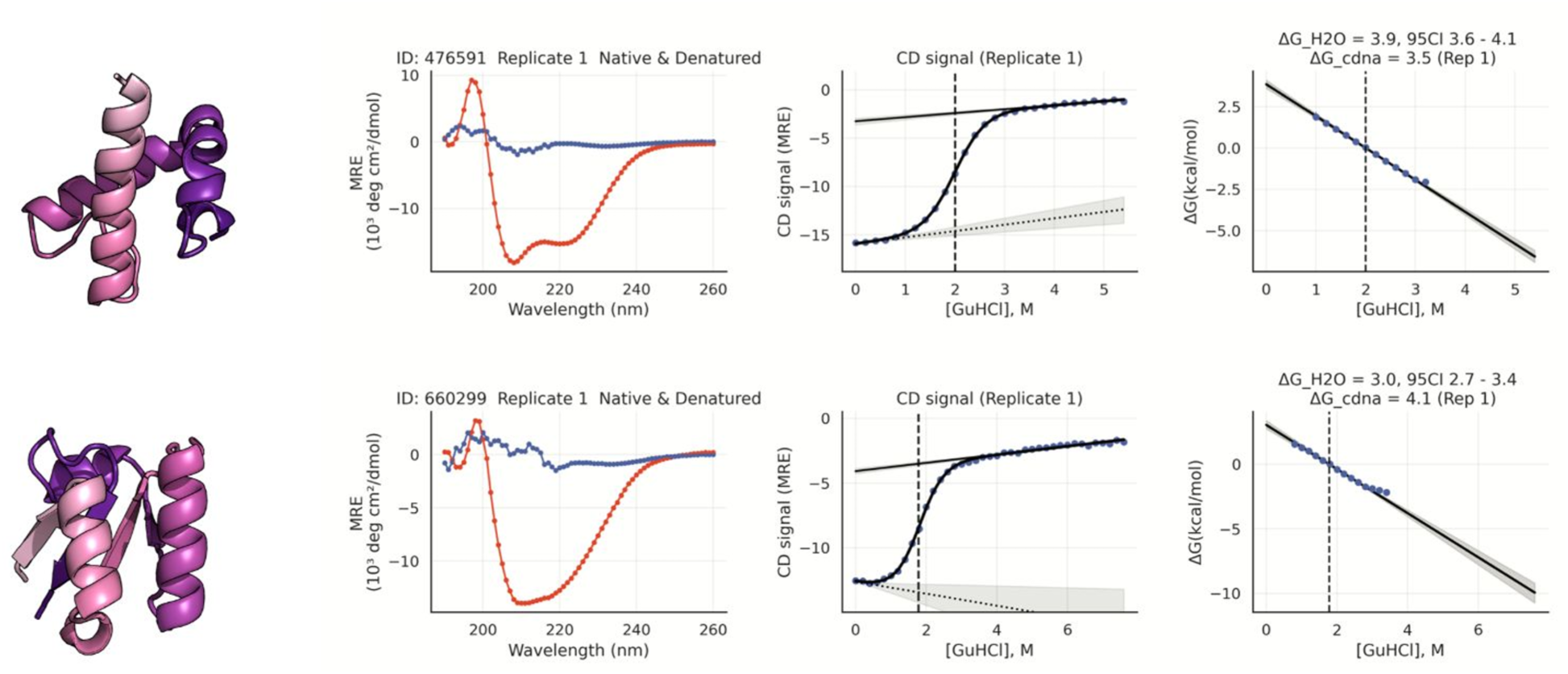
Representative circular dichroism spectroscopy analysis for proteins from the MGnify Stability dataset. **a,** AlphaFold2-predicted structure of a representative protein. **b,** Circular dichroism spectra of the native and denatured states, shown as mean residue ellipticity versus wavelength. **c,** Circular dichroism signal versus guanidine hydrochloride concentration, with unfolding transition and fitted curve. **d,** Free energy of unfolding versus guanidine hydrochloride concentration derived from circular dichroism measurements, with linear fit used to estimate stability.

**Supplementary Figure 3.**
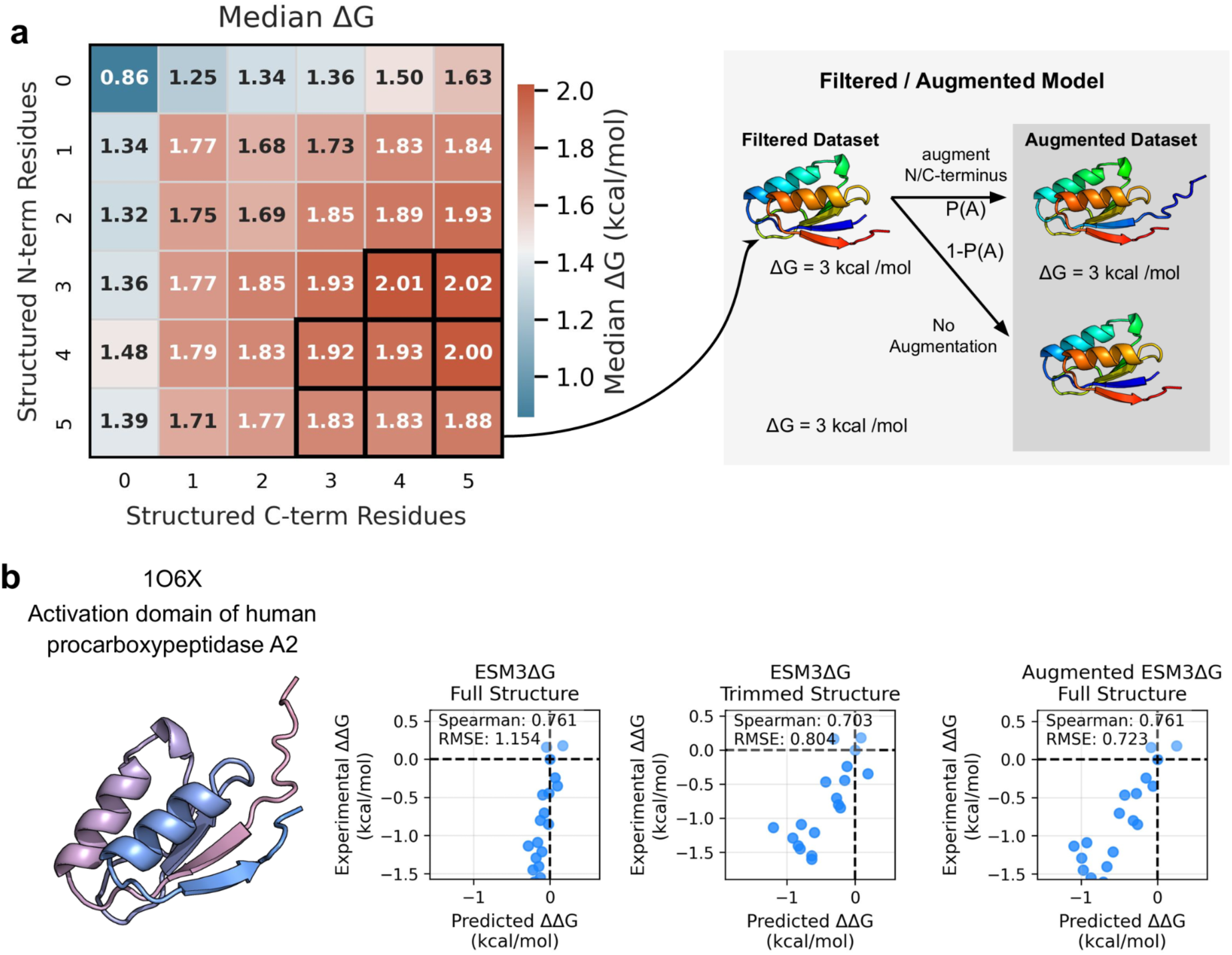
Effect of dataset augmentation on SaProtΔG sensitivity to GS linker insertions. **a,** (Left) Median stability grouped by the number of residues within the first five N-terminal and last five C-terminal positions that adopt secondary structure. (Right) Schematic of dataset augmentation. **b,** Example protein (PDB: 1O6X; activation domain of human procarboxypeptidase A2) illustrating the effect of structural context and dataset augmentation on prediction accuracy. Predicted versus experimental ΔΔG values are shown for ESM3ΔG using full structure (left), trimmed structure with unstructured termini removed (center), and the augmented model on the full structure (right). Spearman correlation and RMSE are shown for each condition.

**Supplementary Figure 4.**
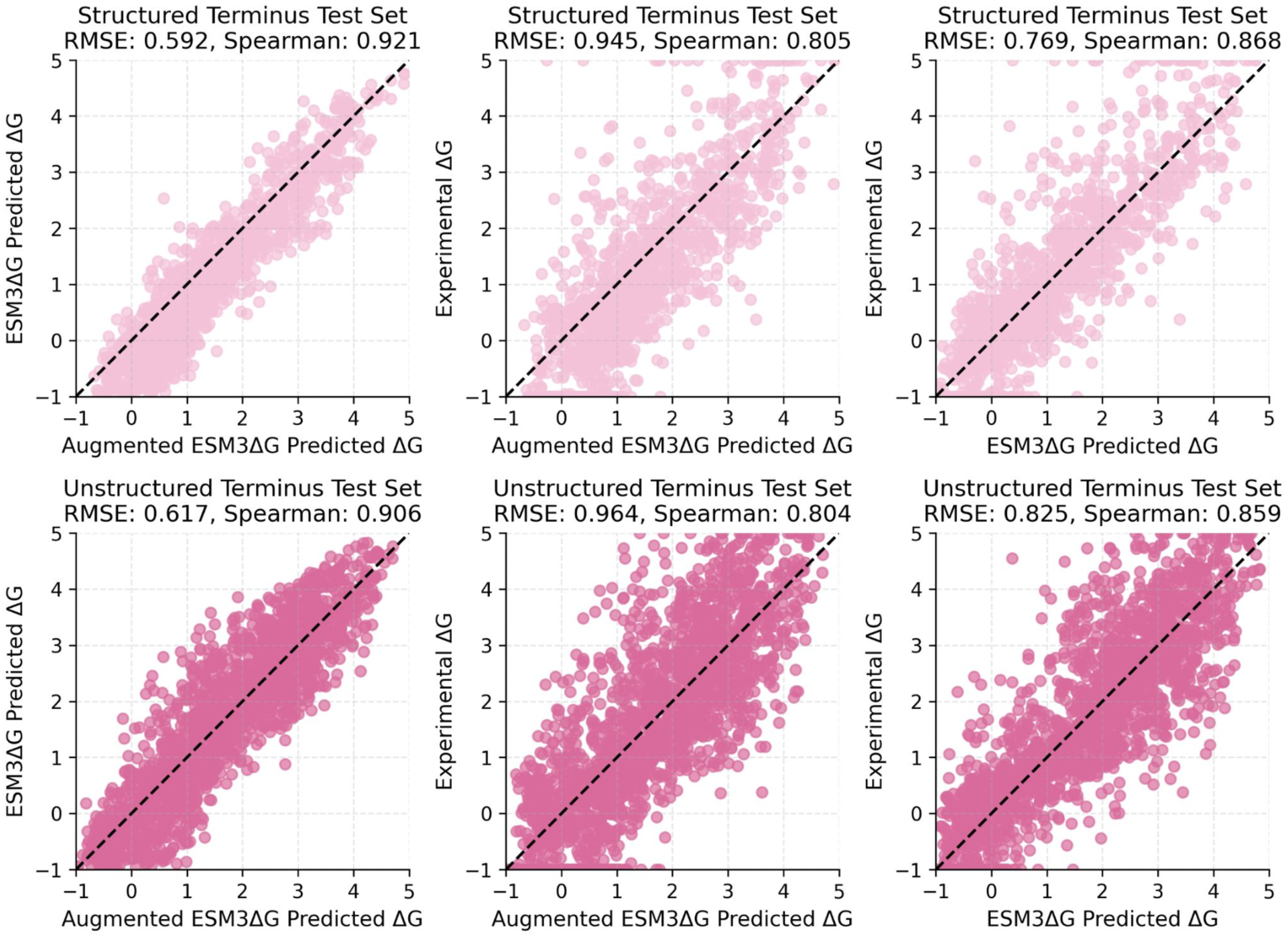
Effect of dataset augmentation on ESM3ΔG prediction accuracy for structured and unstructured termini test sets. ΔG comparisons are shown for structured (top) and unstructured (bottom) terminus test sets. Left, Augmented ESM3ΔG predictions versus ESM3ΔG predictions; center, ESM3ΔG predictions versus experimental ΔG; right, Augmented ESM3ΔG prediction versus experimental ΔG. Each plot shows the RMSE and a black dashed line to indicate y = x.

**Supplementary Figure 5.**
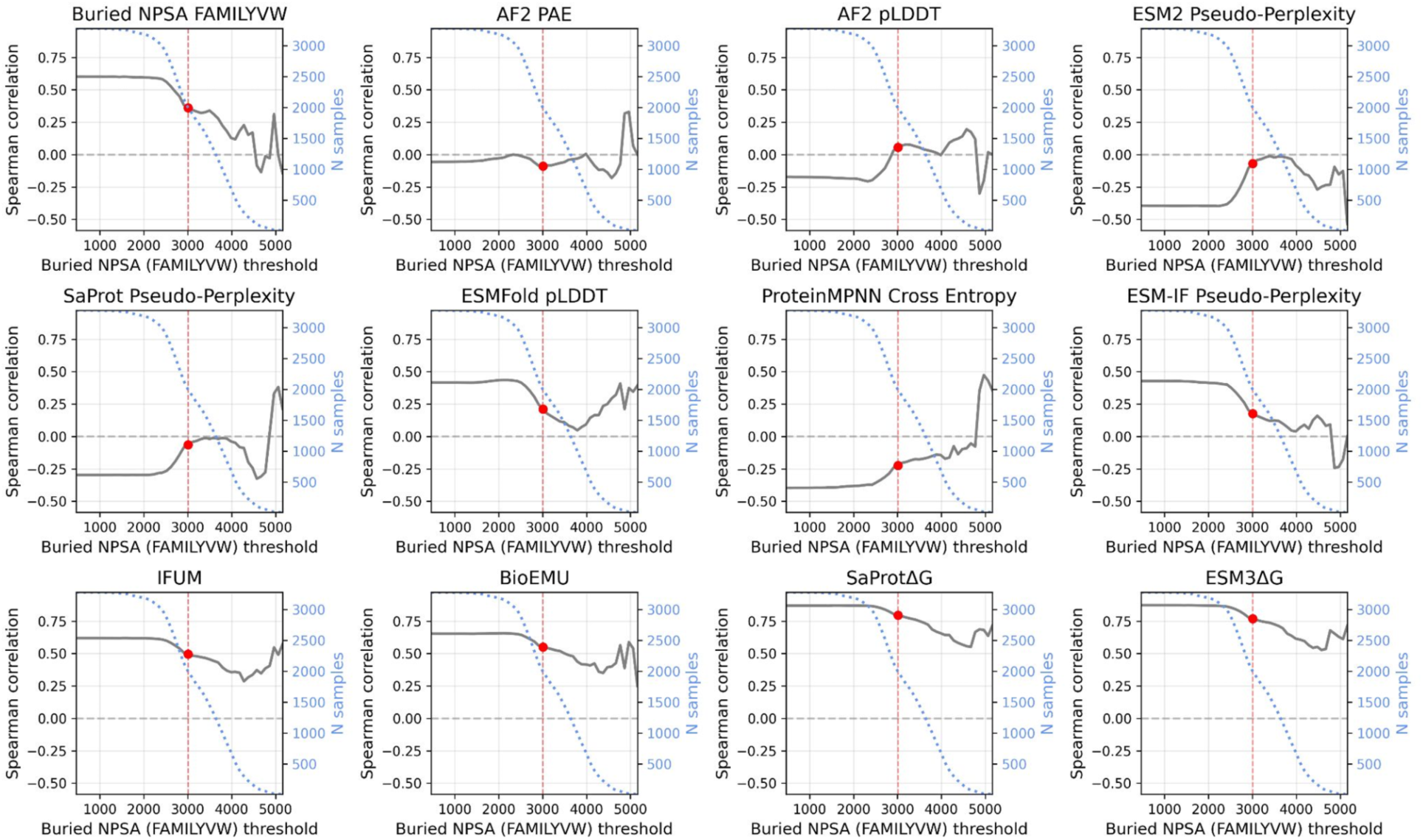
Dependence of prediction accuracy on buried nonpolar surface area (NPSA) thresholding across models. Spearman correlation between predicted and experimental stability is shown as a function of the buried NPSA threshold (FAMILYVW residues) applied to the test set. Each panel corresponds to a different model or scoring function. The gray curve shows correlation (left y-axis) as NPSA thresholds increases. The blue dashed curve indicates the number of sequences remaining at each threshold (right y-axis). Red points and vertical dashed lines mark the 3,000 Å2 threshold.

**Supplementary Figure 6.**
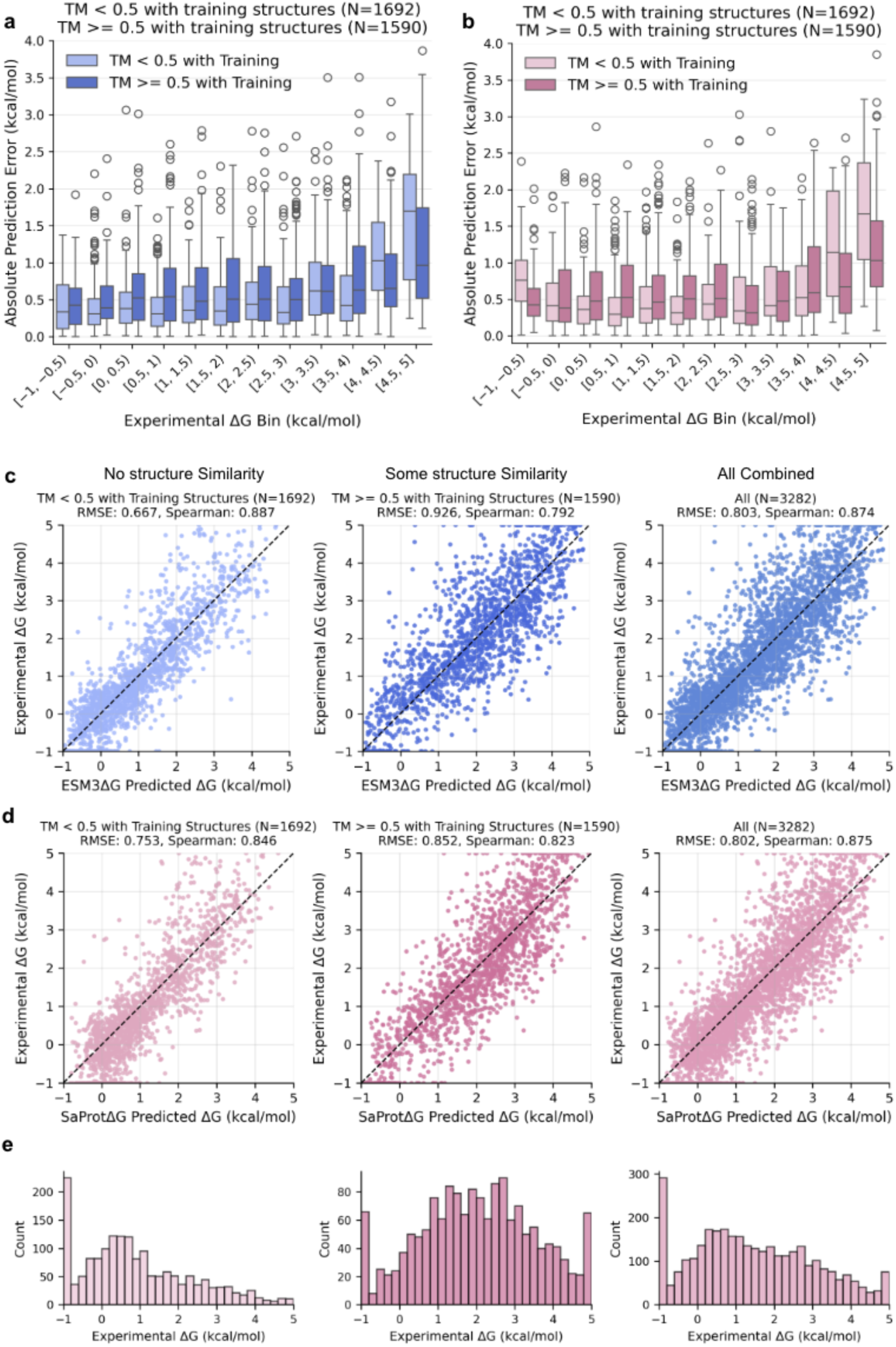
Prediction error as a function of experimental stability and structural similarity to training data. Absolute prediction error (|ΔG_pred − ΔG_exp|) is shown across bins of experimental ΔG for proteins with low structural similarity to training structures (TM < 0.5) and high structural similarity (TM ≥ 0.5). **a,** ESM3ΔG predictions. **b,** SaProtΔG predictions. Box plots indicate median and interquartile range, with whiskers extending to 1.5 times the interquartile range; points denote outliers.

**Supplementary Figure 7.**
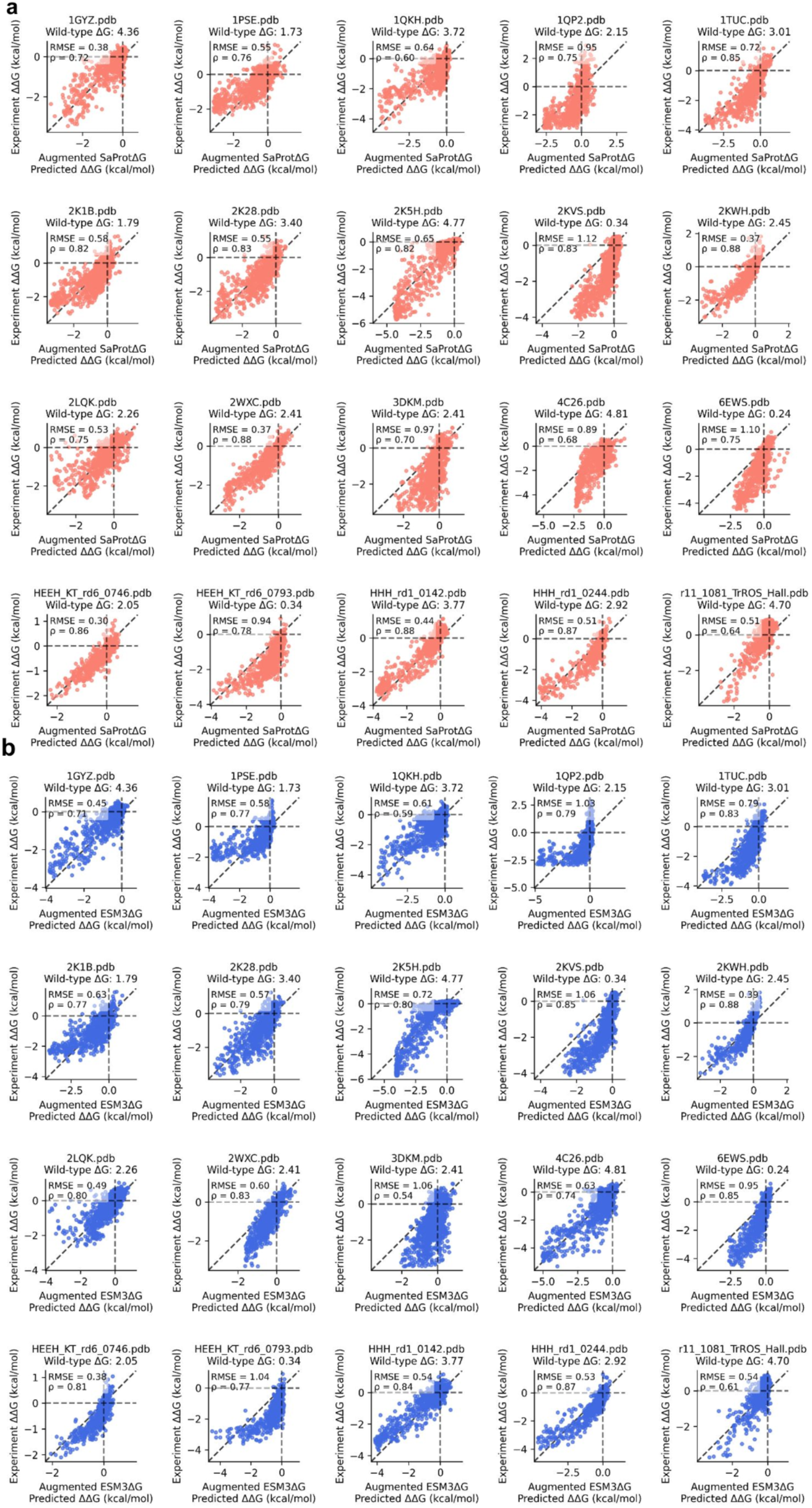
SaProtΔG and ESM3ΔG demonstrate accurate prediction of megascale mutant ΔΔG, with strong agreement to mutational scanning data. **a,** SaProtΔG and **b,** ESM3ΔG predictions of mutational scanning data per PDB. RMSE and Spearman correlation are reported in the plots.

**Supplementary Figure 8.**
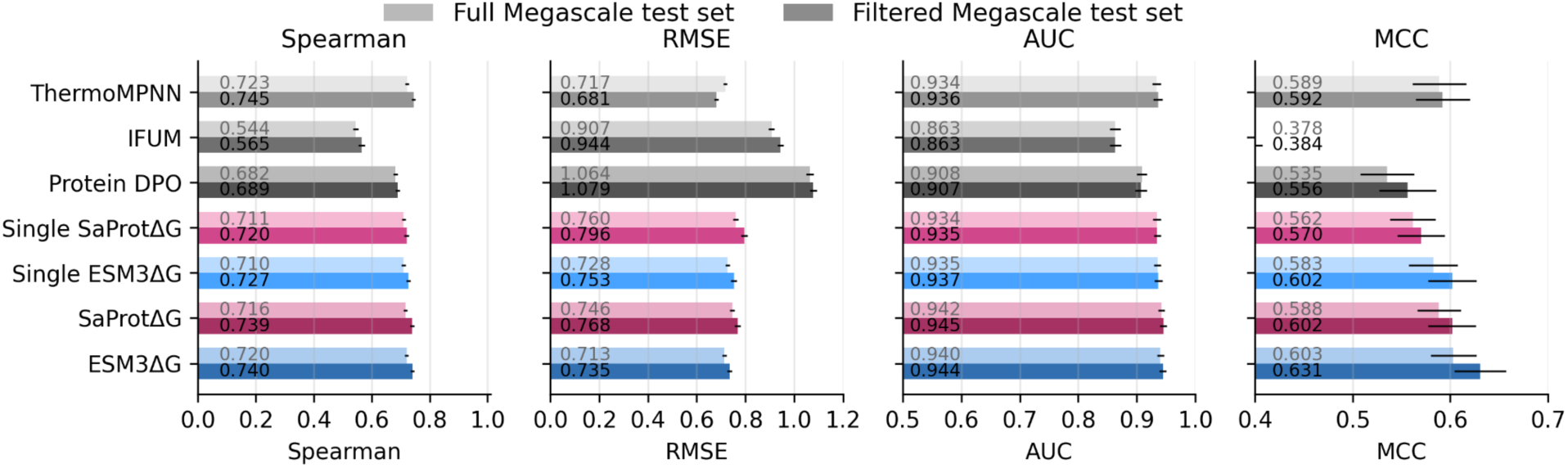
Accuracy on the ThermoMPNN test set after removing wild-type sequences with similarity to the MGnify training set. Model performance is evaluated on the full Megascale test set (light bars) and a filtered subset (dark bars) in which wild-type sequences with sequence identity > 0.3 to at least one protein in the MGnify training data have been removed. Metrics shown are Spearman correlation, root mean squared error (RMSE), area under the ROC curve (AUC), and Matthews correlation coefficient (MCC). Error bars indicate uncertainty across bootstrap resamples.

**Supplementary Figure 9.**
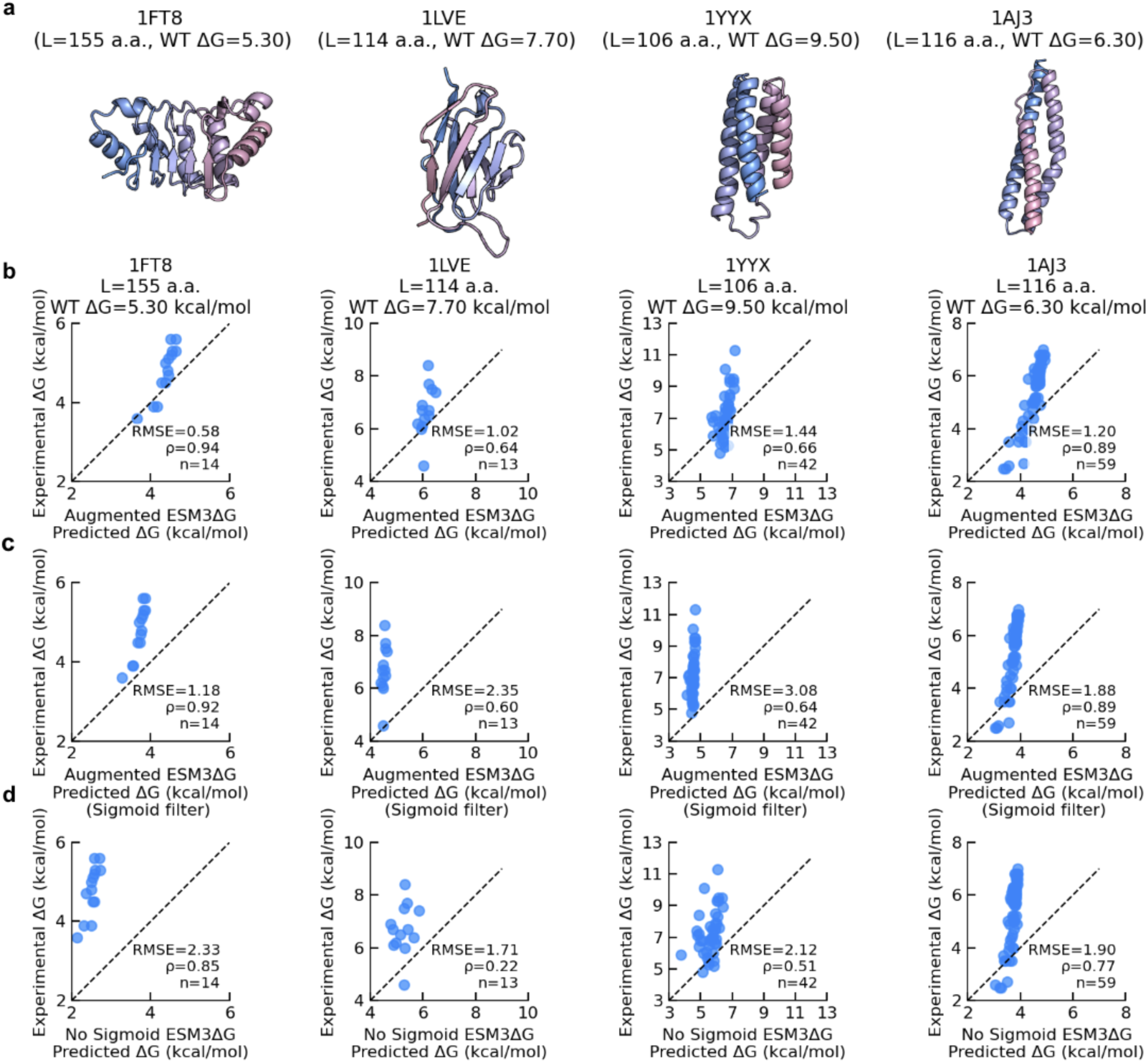
Comparison between sigmoid and non-sigmoid versions of ESM3ΔG. **a,** Structures of 1FT8, 1LVE, 1YYX, and 1AJE. **b,** Augmented ESM3ΔG and **c,** augmented ESM3ΔG with a sigmoid filter. **d,** Non-sigmoid ESM3ΔG evaluated on these proteins, all of which have lengths greater than those in the training dataset (i.e., >100 residues).

**Supplementary Figure 10.**
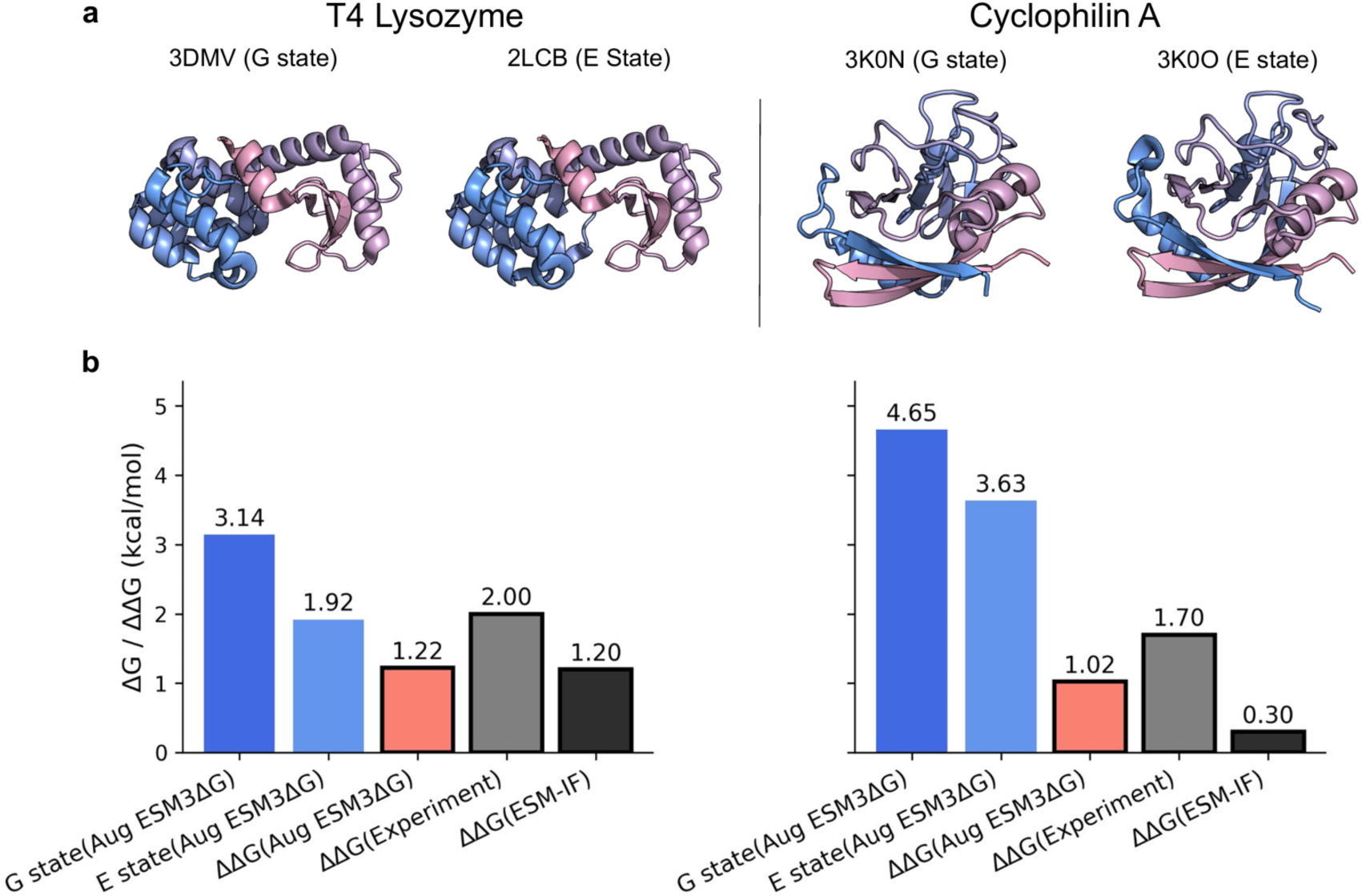
Evaluation of AugmentedESM3ΔG for predicting stability differences between alternative protein conformations. Performance of Augmented ESM3ΔG in predicting absolute folding free energies (ΔG) and the corresponding stability differences (ΔΔG) between distinct conformational states of T4 lysozyme and cyclophilin A, considering both ground (G) and excited (E) states. **a,** Structures of the ground and excited conformations for each protein. (B) Comparison of predicted absolute stabilities and inferred ΔΔG values with experimentally measured ΔΔG, as well as ΔΔG predictions from ESM-IF. In panel **b,** results for T4 lysozyme are shown on the left and for cyclophilin A on the right.

**Supplementary Table 1.**
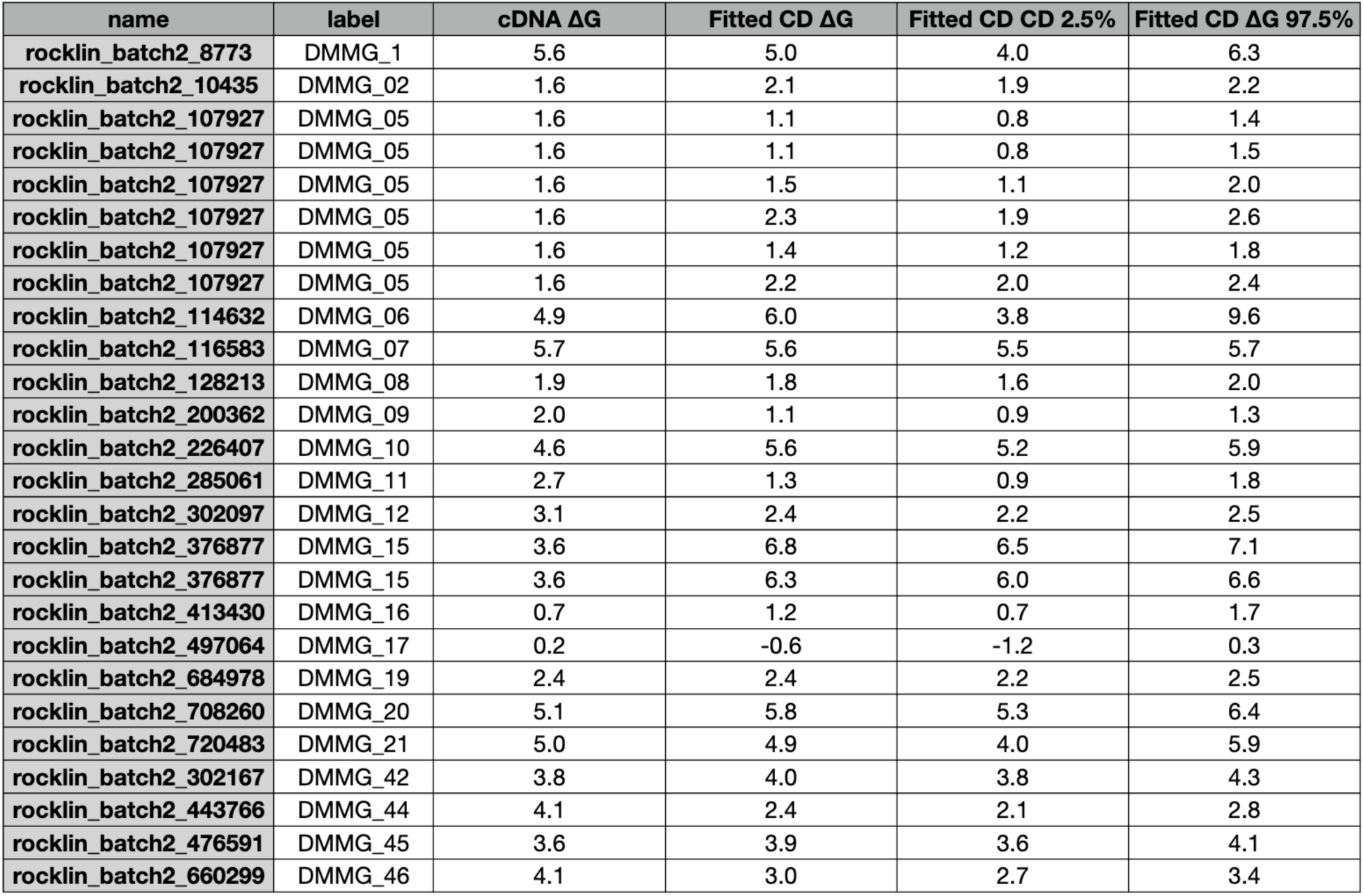
Comparison of CD-measured and cDNA folding stabilities for ThermoMut variants. For each protein variant, CD-measured stability values (Fitted CD ΔG) are compared with cDNA ΔG values. Fitted values are reported as posterior medians, with corresponding 95% credible intervals (2.5% and 97.5% quantiles).

**Supplementary Table 2.**
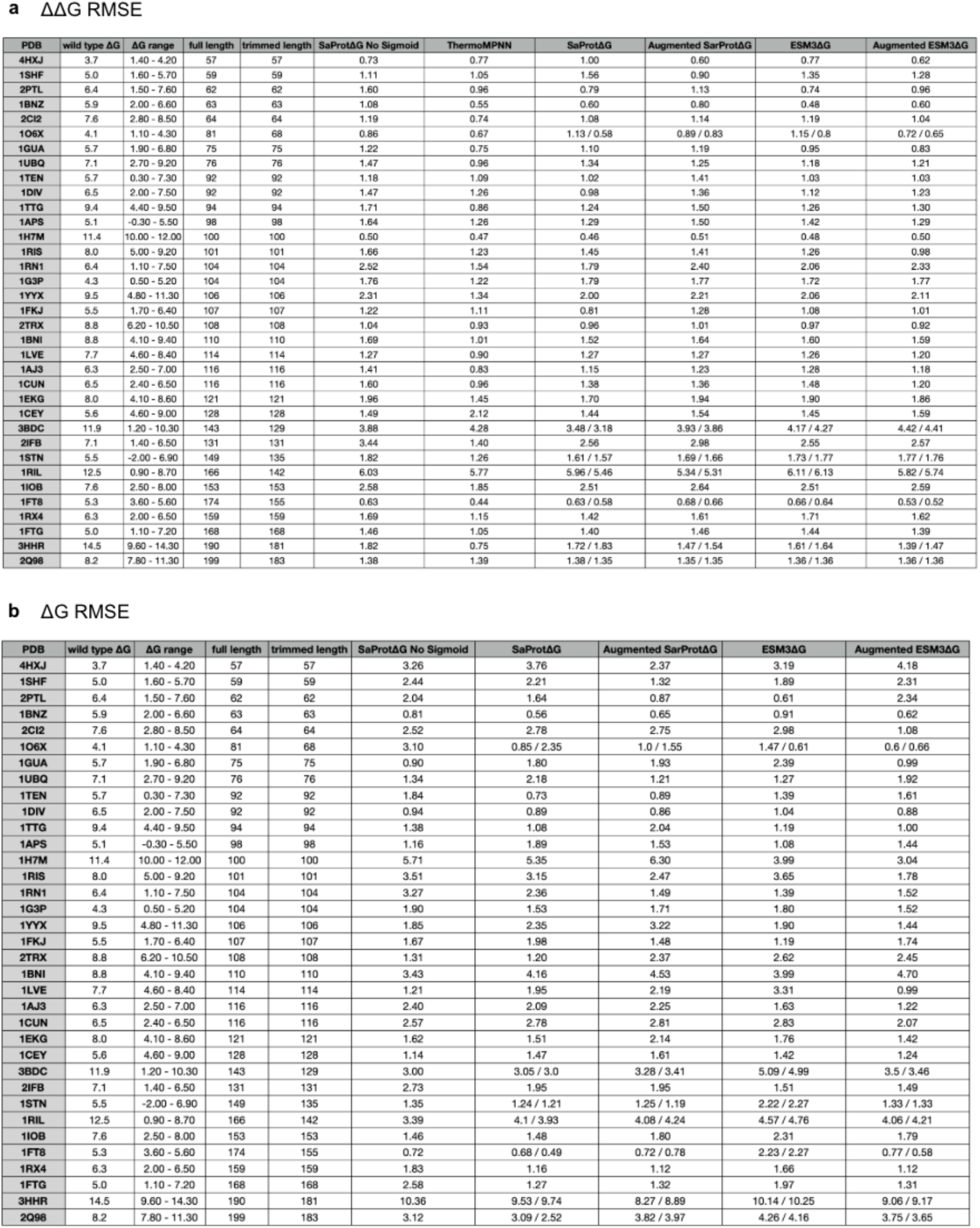
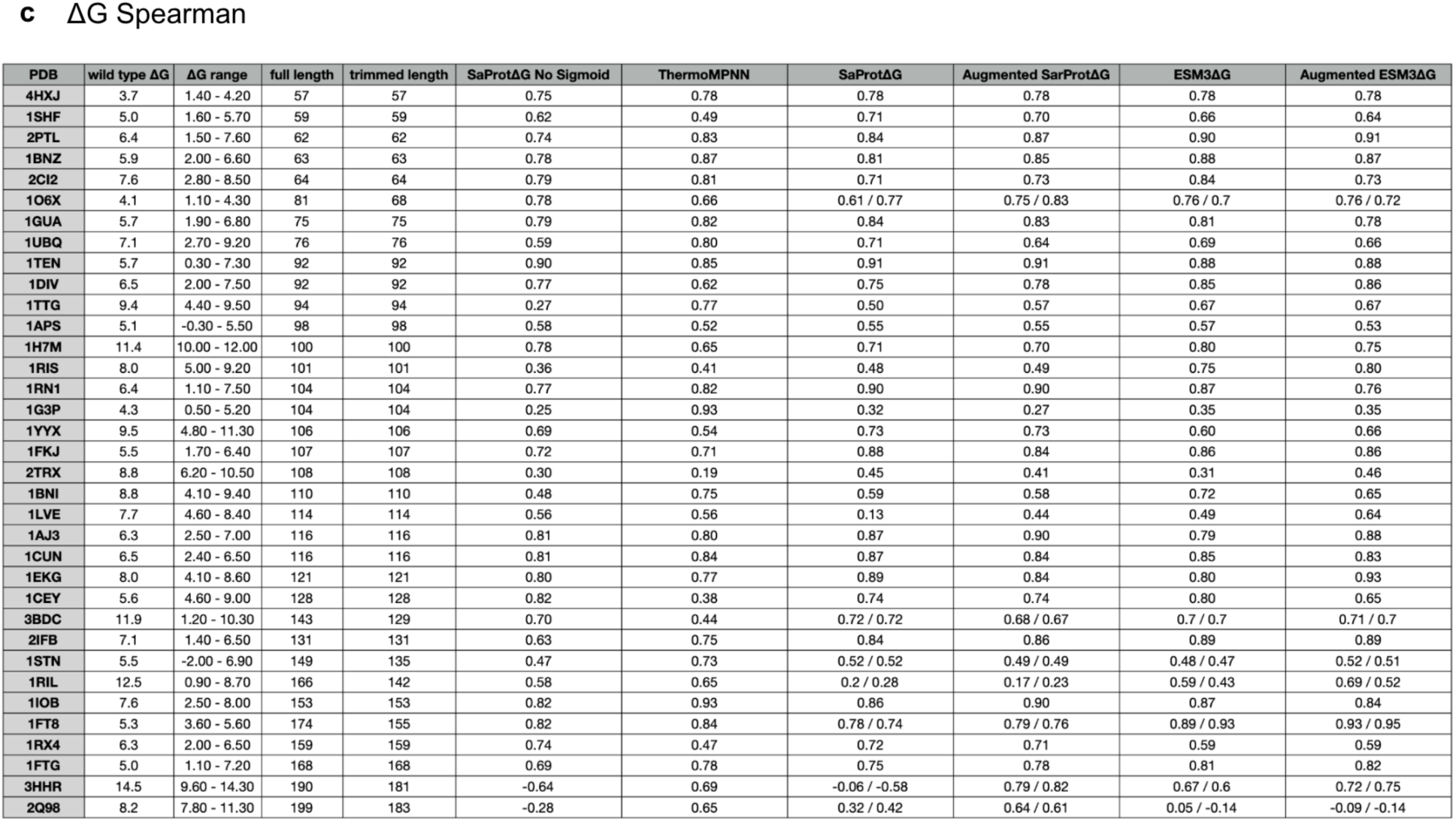
Performance of model predictions on the ThermoMut dataset. **a,** ΔΔG RMSE, **b,** ΔG RMSE, and **c,** Spearman correlation per PDB, comparing ThermoMPNN with SaProtΔG and ESM3ΔG models, with and without augmentation.

## Materials and Methods

### Library design and clustering

The sequences in the MGnify Stability dataset (both wild-type sequences and mutants) were all provided by DeepMind. Briefly, AlphaFold2 structures were determined for all MGnify sequence clusters with 10 or more proteins, then these structures were segmented into putative independent domains by clustering using the Clauset-Newman-Moore method (Clauset et al. 2004). These resulting domains were filtered to remove Cys-containing domains, domains outside the length range of 60-80 amino acids, domains with mean pLDDT < 50, and domains that were a single long helix or otherwise lacked secondary structure. Finally, 730,000 wild-type sequences were randomly chosen from the remaining domains for experimental testing. Single mutants, multiple mutants, and indel mutants were designed from these wild-type sequences using both random sampling and BLOSUM-biased sampling. A selection of AI-designed sequences (not from MGnify) are also included in the experimental dataset but are not analyzed in this study. All MGnify sequences (wildtypes and variants) are labeled “mgnify==True” in the dataset file 230515_K50dG_dmsv4_dmsv5_dmsv7_concat260429.csv.

The full MGnify Stability dataset was experimentally analyzed using cDNA display proteolysis (Tsuboyama et al. 2023), but only a subset of sequences were used for machine learning. To construct the machine learning set, we first used preliminary experimental data quality metrics to exclude low confidence measurements. Based on initial data indicating some highly hydrophobic sequences may appear spuriously stable, we also excluded highly hydrophobic sequences (fraction of FAMILYVW amino acids >= 0.5) from machine learning. All remaining sequences were clustered by MMseqs2 (Steinegger and Söding 2017) at 30% identity and clusters were randomly assigned to training, validation, or test sets. The 30% identity threshold was selected based on empirical analysis showing that homologous proteins at this evolutionary distance still display substantial stability divergence (Fig. 1d), ensuring that predictive performance could not be explained by trivial sequence similarity. Finally, the training set was further filtered using JackHMMER (Finn et al. 2011) to remove training sequences with detectable similarity to the evaluation sets. The final dataset used for machine learning comprised 966,000 sequences (including both wild-type proteins and substitution variants) and was divided into 960,216 training sequences, 3,283 test sequences, and 2,357 validation sequences. No indel variants were used in training. The full machine learning dataset was finalized prior to model training and was not further modified.

To assess potential structural overlap, Foldseek clustering was performed using an easy-cluster TM-score threshold of 0.5. In total, 1,590 unique test structures were found to cluster with at least one training structure. To examine whether this affected performance on the test set, we compared the RMSE distributions for test proteins above and below the TM-score threshold of 0.5. Overall, the RMSE differences between the high- and low-TM-score subsets were minimal, indicating that model performance is not driven by structural similarity or information leakage from the training data (Supplementary Fig. 6).

After model training, we re-calibrated the original experimental data to incorporate additional mutant and indel data (the “v7” library). This final recalibrated data is shared on Zenodo (https://forms.gle/4ZnXZSnTBvaykkAi9) in the file 230515_K50dG_dmsv4_dmsv5_dmsv7_concat260429.csv. This recalibration resulted in modified stability estimates for the sequences used in training, which typically changed by 0.1-0.2 kcal/mol. The final recalibrated deltaG estimates are given in the “deltaG” column; the values used for training are given in the “train_dG_from_earlier_processing” column.

### cDNA display proteolysis

We followed the protocol as described (Tsuboyama et al. 2023), and calibrated ‘effective’ protease concentrations using the previous dataset (Tsuboyama et al. 2023). As in (Tsuboyama et al. 2023), the ∼2,000 overlapping sequences between each library were used to calibrate effective protease concentrations for each of the 48 experimental conditions per experiment (24 trypsin conditions and 24 chymotrypsin conditions). The consistency between experiments for these overlapping sequences shown in Supplementary Figure 1 B-C is after calibration. Scripts to process the original NGS counts data into ΔG estimates are provided on Zenodo (https://forms.gle/4ZnXZSnTBvaykkAi9).

### Circular dichroism

All far-ultraviolet CD measurements were done on a Jasco J-1500-150 (Supplementary Fig. 2, Supplementary Table 2). Wavelength scans were measured from 260 to 190 nm at 25°C, at a scanning speed of 50 nm/min, 8 sec. D.I.T., with 2 accumulations. Wavelength scans were performed immediately prior to and after a chemical denaturation run with 0.03 mg/ml protein in PBS buffer sans NaCl (10 mM NaPO4 and 1.8 mM KPO4 pH 7.4) with a 1 cm path-length cuvette. For the post melt wavelength scan, the guanidinium hydrochloride (GuHCl) concentration varied depending on the individual protein. Chemical denaturation experiments with GuHCl were performed using an automatic titrator with a protein concentration of 0.03 mg/ml and a 1 cm path-length cuvette with stir bar. The GuHCl concentration was determined by refractive index in PBS buffer using a Brix digital refractometer (Fisherbrand HDR-P1). The denaturation process monitored dichroism signal at 222 nm in steps of 0.1 M or 0.2 M GuHCl with 75 seconds mixing time for each step, at 25°C. All protein concentrations were determined by Qubit protein assay (ThermoFisher Scientific Q33212). Chemical denaturation curves were fitted using a Bayesian nonlinear regression model to a two-state unfolding model with six-parameters: the folding free energy, m-value, and linear pre- and post-transition baselines with individual slope and intercepts (Santoro and Bolen 1988). Raw CD data and CD processing scripts are provided on GitHub (https://github.com/yehlincho/absolute-stability-predictor).

### Biophysical feature analysis (Fig. 2A)

Biophysical properties for the MGnify test set (including buried nonpolar surface area in Fig. 2A) were computed using the featurization pipeline available at https://github.com/Rocklin-Lab/mhdx_analysis/tree/main/scripts/feature_extraction as previously used in (Ferrari et al. 2025). “Buried NPSA FAMILYVW” refers to buried nonpolar surface area from the subset of nonpolar amino acids Phe, Ala, Met, Ile, Leu, Tyr, Val, and Trp.

### Megascale point mutation benchmark (Fig. 3B-C)

This benchmark included 28,172 substitution mutants from 28 wild-type proteins from the original Megascale stability dataset (Tsuboyama et al. 2023), as originally curated for ThermoMPNN benchmarking ((Tsuboyama et al. 2023; Dieckhaus et al. 2024).

### MGnify insertion and deletion benchmark (Fig. 3D)

From all insertions and deletions included in the MGnify Stability Dataset, we filtered a subset of the highest quality measurements to use for benchmarking. Entries were retained only if the 95% confidence interval of the free energy estimate (ΔG) was smaller than 0.4 kcal/mol, fitting fitting errors for both trypsin and chymotrypsin conditions were required to be below 0.17, and the difference between independent trypsin and chymotrypsin measurements (|ΔG_trypsin − ΔG_chymotrypsin|) was below 1.5. Only insertions and deletions satisfying all of these criteria were retained for downstream evaluation.

### Megascale stabilizing insertions and deletions benchmark (Fig. 3E)

This set of 91 stabilizing mutants (ΔΔG_unfold_ > 1) stabilizing insertions and deletions was originally measured in the Megascale stability dataset (Tsuboyama et al. 2023), as curated by (Gutierrez and Rocklin 2024).

### The S1724 benchmark of ΔG data from larger proteins assayed using traditional methods (Fig. 4)

This set was constructed by manually selecting subsets of mutants from ThermoMutDB (Xavier et al. 2021; Zwanzig 1997)). Mutant sets were included in S1724 if (1) at least 10 variants were characterized in a single publication from a single lab, (2) folding of the wild-type protein was shown to be reversible (3) folding measurements were done with chemical denaturation using either urea or guanidine chloride and could be fit using a two-state model, and (4) folding measurements were conducted in buffers between pH 5 and 8 at temperatures between 15°C and 30°C. We verified reversible, two-state folding by inspecting the original experimental publications compiled into ThermoMutDB. Some measurements are linear extrapolation of equilibrium experiments and others are linear extrapolation of kinetic fits.

The original ThermoMutDB database compiles ΔΔG measurements, not ΔG. Some of these ΔΔG measurements are extrapolated back to pure buffer conditions with no denaturant, whereas others indicate ΔΔG at a specific denaturant concentration (e.g. ΔΔG at the wild-type midpoint concentration). In some cases, ThermoMutDB averages equilibrium ΔΔG estimates with kinetic ΔΔG estimates to produce a single ΔΔG entry. To compile the ΔG estimates in S1724, we typically started from ThermoMutDB ΔΔG (regardless of the method of computation) and added these to wild-type ΔG values typically from the original reference used by ThermoMutDB. We comprehensively inspected ThermoMutDB ΔΔG values for consistency with the original publications and corrected all errors based on the original publications. A comprehensive list of our corrections and changes to ThermoMutDB is available on Zenodo (https://forms.gle/4ZnXZSnTBvaykkAi9).

### Benchmarking correlations with optimal growth temperature (Fig. 5)

Structural domain annotations were obtained from the TED (The Encyclopedia of Domains) dataset (Lau et al. 2024). Entries were restricted to single-domain proteins ≤ 200 residues and pLDDT ≥ 70. Growth temperatures were assigned by mapping proteome_id-derived taxonomy identifiers to an external dataset (Martin Karl Magnus Engqvist 2018) via tax_id. Entries lacking UniProt mappings, duplicates, or sequences with conflicting temperature annotations were removed. Within each CATH classification, sequences deviating by >20% from the median domain length were excluded. To reduce imbalance, CATH groups with >5,000 sequences were downsampled to 5,000 using a uniform temperature distribution. CATH groups with <1,000 sequences or <100 sequences above 50 °C were excluded. To place the ProteinMPNN cross-entropy (CE) and AlphaFold2 pLDDT metrics on a comparable scale with the predicted absolute stability (kcal/mol) estimated using Augmented ESM3ΔG, we applied a quantile-matching transformation. First, the empirical distribution of Augmented ESM3ΔG predicted ΔG values across the full dataset was computed and sorted from high stability to low stability to define a reference distribution. For each metric (CE and pLDDT), percentile ranks were calculated using their empirical cumulative distributions. These percentile values were then mapped to the corresponding quantiles of the Augmented ESM3ΔG reference distribution, thereby transforming each metric to follow the same marginal distribution as the predicted stability. Because lower CE values correspond to stronger sequence compatibility, CE values were sign-inverted prior to ranking to preserve the directionality of the relationship. This non-parametric transformation preserves the relative ordering of scores while aligning their distributions with that of the predicted stability estimates, enabling direct comparison and integration of the different metrics.

### Benchmarking on de novo designed proteins (Fig. 6A-D)

Rosetta designed proteins with annotated success/failure were taken from (Garcia et al. 2026), a compilation of 12 previously published studies (2012–2021). De novo designs in Fig. 4c (5,000 backbones × 4 design tools) measured for folding stability by cDNA display proteolysis were taken from (Cho et al. 2025).

### Benchmarking on de novo designed binders (Fig. 6H-I)

Binder and non-binder sets were designed using RFdiffusion on five targets (Watson et al. 2023). For RFdiffusion, we further restricted the analysis to monomeric proteins. We also included EGFR binders from the Adaptyv Bio competition (Cotet et al. 2025), where we restricted the dataset to *de novo* proteins, excluding any antibody or VHH sequences, resulting in 302 designs.

All AUC values reported in this paper refer to the area under the receiver operating characteristic curve (ROC AUC), computed for binary classification tasks (e.g., folded vs. unfolded, binder vs. non-binder, or experimental success vs. failure of de novo designed proteins).

### Baseline Model Stability Prediction

To benchmark our models against the existing literature (Fig. 2), we computed predictions from a range of biophysical descriptors, zero-shot model scores, and externally trained models. These are organized into four comparison tiers below.

- **Zero-shot features** Biophysical properties (longest loop length, sequence length, radius of gyration, nonpolar residue count, buried nonpolar surface area, and “Buried NPSA FAMILYVW”) for the MGnify test set were computed using the featurization pipeline described in the Biophysical feature analysis section above (Ferrari et al. 2025). These features require no machine learning inference and serve as purely sequence- and structure-derived stability proxies.
- **Zero-shot confidence**

– Per-residue pLDDT and predicted aligned error (PAE) from AlphaFold2 (Jumper et al. 2021) were used directly as stability proxies, as higher structural prediction confidence has been linked to thermostability (Cho et al. 2025). ESMFold (Lin et al. 2023) pLDDT scores were computed from ESMFold-predicted structures of each MGnify test sequence.
– ESM2 pseudo-perplexity was computed using ESM2 (facebook/esm2_t33_650M_UR50D; (Lin et al. 2023)) following the pseudo-log-likelihood approach: each residue position was masked in turn, the masked-language-model loss was evaluated at that position, and the per-position losses were averaged. Pseudo-perplexity is defined as the exponent of this mean loss; negative pseudo-perplexity (higher probability = lower perplexity = greater estimated fitness) was used for correlation analysis.
– SaProt pseudo-perplexity was computed using SaProt_650M_AF2 (Su et al. 2023), with the distinction that SaProt’s input interleaves amino acid tokens with Foldseek structure tokens derived from the AlphaFold2-predicted structure, so structural information is incorporated into the scoring.
– ESM-IF ((Hsu, Nisonoff, et al. 2022) esm_if1_gvp4_t16_142M_UR50) is a GVP-based inverse folding model conditioned on backbone coordinates. For each test protein, the sum of per-position log-probabilities of the native amino acid was computed. This raw score was then linearly calibrated to kcal/mol units (ΔG = 0.104 × ΣlogP(aa_i) + 0.616) using a previously fitted linear regression (Cagiada et al. 2025). ProteinMPNN (Dauparas et al. 2022) cross-entropy was computed using the v_48_010 model via ColabDesign (Moffat et al. 2022); the per-residue cross-entropy of the native sequence given the structure was averaged as a stability proxy.
- **Megascale (external fine-tuned models)**

– ProteinDPO (Widatalla et al. 2024): ProteinDPO fine-tunes ESM-IF1, a structure-conditioned inverse folding language model, using Direct Preference Optimization on stability-ranked sequence pairs from the Megascale dataset. The DPO objective trains the model to assign higher likelihood to more thermostable sequence variants over less stable counterparts from the same backbone. At inference, the stability score for a test sequence is the log-likelihood assigned by the DPO-fine-tuned ESM-IF1 model given the protein backbone structure (https://github.com/evo-design/protein-dpo).
– IFUM (Lee et al. 2026): IFUM is a transformer-based deep neural network that jointly predicts ΔG and the equilibrium ensemble of folded and unfolded states. The model takes as input sequence embeddings from ProtT5, structural embeddings from ESM-IF1, and ESMFold-predicted distograms; these are processed by a modified Evoformer to output per-residue free energy contributions that are summed to yield the total ΔG. Predictions were obtained by running the published IFUM inference script on AlphaFold2-predicted structures of each test sequence (https://github.com/HParklab/IFUM).
- **Megascale+ (external fine-tuned model)**

– BioEMU 1.2 (Lewis et al. 2025) is a generative diffusion model trained to sample Boltzmann-distributed protein structural ensembles through three stages: pretraining on AlphaFold Database structures, fine-tuning on ∼145 ms of aggregate MD trajectories, and Property Prediction Fine-Tuning (PPFT) that aligns the sampled ensemble to experimental ΔG labels. BioEMU 1.2 was trained on the Megascale cDNA display proteolysis dataset plus additional unpublished cDNA display proteolysis measurements. Predictions were obtained using the updated BioEMU model weights (https://github.com/microsoft/bioemu).

### Fine-tuning Absolute Stability Models

Large protein models can predict stability in zero-shot settings (Notin et al. 2023), and incorporating structural information further enhances accuracy (Cho et al. 2024). Structural features such as buried surface area and solvent-accessible surface area have been linked to protein stability, motivating the use of structure-informed architectures.

To establish a simple baseline and evaluate how much predictive signal can be learned without large pre-trained models, we trained a lightweight model from scratch. In this baseline, amino acid sequence tokens were combined with structure tokens derived using Foldseek. Both sequence and structure tokens were represented using one-hot encoding and concatenated into a single feature representation.

These features were used to train a small multi-layer perceptron (MLP) to predict protein stability (ΔG). We refer to this model as Sequence-Structure MLP, which provides a minimal architecture capturing sequence structure information and serves as a baseline for comparison with large pre-trained models.

Building on this baseline, we benchmarked and fine-tuned four protein models:

- ProteinMPNN (Dauparas et al. 2022)(structure-to-sequence)

– We finetuned ProteinMPNN end-to-end for protein stability prediction, following a similar strategy to ThermoMPNN (Dieckhaus et al. 2024) but applied to full sequences rather than point mutations. Residue-level features are formed by concatenating hidden representations from the top two encoder layers with sequence embeddings; these are processed by a lightweight attention-based head to produce per-residue ΔG estimates. Global stability is obtained by averaging per-residue predictions, and training minimizes the error between predicted and target ΔG.
- AlphaFold2 (AF2) (Jumper et al. 2021)(sequence-to-structure)

– We fine-tune AlphaFold2 by introducing an additional per-residue ΔG prediction head on top of the single representation. The head comprises a multi-block attention module (4 blocks, 16 heads, key dimension 16), followed by a transition layer with a 2× channel expansion and a final linear projection to produce per-residue stability predictions.
– During fine-tuning, template information and distillation data are disabled. The model is optimized using a multi-task objective that combines stability and structure-related losses, with weights: ΔG (2.4), masked MSA (2.0), structure module (1.0), distogram (0.3), PAE (0.1), experimentally resolved (0.01), and pLDDT (0).
- ESM-3 (Hayes et al. 2025) (multimodal sequence/structure-to-sequence/structure)
- SaProt (Su et al. 2023) (multimodal sequence/structure-to-sequence/structure)

AlphaFold2 takes only a sequence and multiple sequence alignment (MSA) as input, while ProteinMPNN, SaProt, and ESM-3 take both a sequence and an AF2-predicted structure. For tokenization, ESM-3 encodes structures into 4,096 tokens, while SaProt uses 20 structure tokens.

We employed two fine-tuning strategies:

1. Full fine-tuning with an MLP head: Model weights were unfrozen and a stability prediction head was added, trained to predict per-residue ΔG values whose average is regressed against experimental ΔG.
2. Low-Rank Adaptation (LoRA): To reduce the number of trainable parameters while retaining pre-trained knowledge, we applied LoRA (Hu et al. 2021) to SaProt and ESM-3. LoRA introduces trainable low-rank matrices:

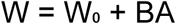

where W₀ ∈ ℝᵈˣᵈ is the pre-trained weight matrix, and B ∈ ℝᵈˣʳ and A ∈ ℝʳˣᵈ are learnable low-rank matrices with r ≪ d.

– We fine-tuned esm3-sm-open-v1, the smallest model in the ESM3 family with 1.4B parameters. LoRA was applied to the query projection and output projection layers of each attention block (target modules: “layernorm_qkv.1” and “out_proj”) with rank r = 4 and dropout = 0.15, requiring updates to only 0.295% of parameters (4.14M out of 1.4B). On top of the frozen (plus LoRA-augmented) ESM3 backbone, a stability head was added: a two-layer MLP consisting of Linear(1536 → 1536) → LayerNorm → ReLU → Linear(1536 → 1), producing per-residue ΔG logits. The per-residue predictions were masked to exclude the [CLS] and [EOS] special tokens, then averaged to produce the global ΔG estimate.
– We fine-tuned SaProt_650M_AF2 using LoRA applied to the query, key, value, and feed-forward projection layers of each transformer block (target modules: “query”, “key”, “value”, “intermediate.dense”, “output.dense”), with rank r = 4 and dropout = 0.15, updating 4.7M out of 656M parameters (0.72%). The stability head is a two-layer MLP: Linear(1280 → 1280) → ReLU → Linear(1280 → 1). Per-residue predictions were masked to exclude padding and special tokens, then averaged to produce the global

### Fine-Tuning with a Sigmoid-Corrected Stability Head

An ideal absolute stability predictor should (i) generalize to experimental baselines that measure folding stability on the real kcal/mol scale, and (ii) predict both wild-type and mutant stabilities, thereby enabling ΔΔG calculation as:

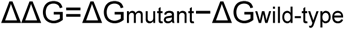

However, cDNA display measurements introduce limitations when assessing proteins at the extremes of stability. In particular, values outside the range of –1 to 5 kcal/mol are often capped or unreliable due to experimental constraints. To address this, We fine-tuned the base models using LoRA and appended a single stability prediction head, a two-layer MLP applied per residue, followed by a piecewise sigmoid correction layer.

### Range-Corrected Activation Function

To accommodate the capped cDNA range while allowing extrapolation, we applied a piecewise activation function with sigmoid tails and a linear middle regime:

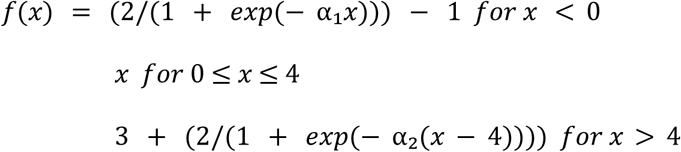

Here, α1 and α2 are trainable slope parameters, allowing the model to learn independent scaling factors and generalize to stability values outside the –1 to 5 kcal/mol training range.

### Loss Function with ΔΔG Regularization

During training, we introduced an additional ΔΔG loss term to explicitly improve the model’s ability to capture mutation effects on stability. The combined mean squared error (MSE) loss was defined as:

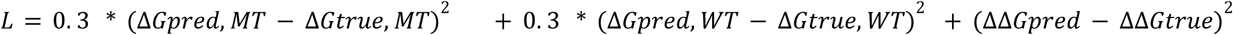

This formulation jointly penalizes errors in wild-type stability, mutant stability, and ΔΔG, ensuring robust predictions across use cases.

At inference time, we activate the sigmoid correction layer for cDNA-based datasets to account for measurement bias. For non-cDNA datasets (e.g., calorimetry), we bypass this layer, under the assumption that the model has already learned to map stabilities to the real kcal/mol scale during training.

### Filter and Augment Terminus Regions to Further Reduce cDNA Display Measurement Bias

To mitigate terminus bias, we first applied a filtering step, excluding sequences with more than two unstructured residues at either terminus. Unstructured residues were identified by scanning the last six residues at both the N- and C-termini, reducing the training dataset to 456,000 sequences. During training, terminus augmentation was applied with 50% probability: a random length of 1–15 residues was selected at each terminus and replaced with randomly sampled amino acids. This strategy reduced the model’s sensitivity to terminus artifacts, encouraging learning of stability determinants from the structured core rather than assay-specific biases at the termini.

